# Evolution of Plant Niche Construction Traits in Biogeomorphic Landscapes

**DOI:** 10.1101/2021.07.18.452868

**Authors:** Xiaoli Dong

## Abstract

By virtue of niche construction traits, plants play a significant role in shaping landscapes. The resultant outcome is a change in the selective environment, which influences the evolution of these same plants. So far almost all biogeomorphic models have assumed that niche construction traits are invariant in time. On the other hand, niche construction studies have assumed that independent abiotic changes are either nonexistent or are simply linear. Here, I considered the concomitant evolution of plant niche construction traits during landscape development. I constructed a geo-evolutionary model that couples a population genetic module with a landscape development module. Allowing plants to evolve always results in landforms different from those that appear when evolution is not accounted for. The topographic difference between cases with and without evolution ranges from a small difference in the steady-state topography, to drastic differences in landforms. The amount of difference is contingent upon forms of landscape development and the strength of geo-evolutionary coupling. Allowing the landscape to develop while evolution occurs changes evolutionary trajectories for niche construction traits. The landscape can even develop spatial structures that suppress selection. Model results clearly support the need to integrate niche construction theory and biogeomorphology to better understand both.

## 1. Introduction

Landscape morphology is shaped by abiotic processes such as uplift, deformation, breakdown, and chemical weathering of bedrock and the erosion, transport and deposition of sediments. These processes occur whether life is present or not; however, living organisms can influence these abiotic processes, and leave distinct topographic signatures at various scales (Dietrich and Perron 2006). Recognition of this role of organisms, especially vegetation, has led to the emergence of the new field of biogeomorphology. Biogeomorphic feedbacks between plants, water or air flow, and sediment transport are a major determinant of large-scale landscape development in tidal marsh landscapes (Kirwan and Murray 2007; Temmerman et al. 2007), alluvial floodplain rivers (Murray and Paola 2003; Gurnell and Petts 2006; Tal and Paola 2007), fluvial hillslopes (Collins 2004; Istanbulluoglu and Bras 2005), and aeolian dune landscapes (Baas and Nield 2007; Hacker et al. 2019). In all these systems, the physical structure of plants imposes friction on flows, and thereby modifies erosion and sedimentation, thereby altering local patterns of elevation by which biogeomorphic change can be reckoned. This environmental change feeds back to affect plants. Plant modification of landscapes can be species-specific, and is conferred by the unique traits of the plant, such as structural traits (e.g., density, length, flexibility of plant shoots) (Bouma et al. 2013), life-history traits (e.g., colonization rate) (Schwarz et al. 2018), and dispersal strategies (Reijers et al. 2019).

Outcomes of these environmental modifications by plants can be neutral, beneficial, or harmful to the plants themselves, but in any case, they can alter the selective environment of these same plants (Matthews et al. 2014). Niche construction theory posits that organism-mediated modifications of the environment can change selection pressures and influence the evolutionary trajectories of natural populations (Laland, Odling-Smee, and Feldman 1999) (Fig. 1). Effects of landscape morphology and physical processes on biology tend to operate over the longer timescale of the development of the landscape pattern. Thus, these effects are not often as obvious as the effects of biology on physical processes, which can occur more quickly (Murray et al. 2008). As a result, research in biogeomorphology to date has almost exclusively focused on the effect of biological processes on landforms (e.g., Bouma et al. 2013; Schwarz et al. 2018), and has considered plants and their niche-constructing traits to be invariant in time, despite ample evidence to the contrary (Van Hulzen, Van Soelen, and Bouma 2007; Reijers et al. 2020). Feedbacks between biological evolution and landscape development have been rarely investigated beyond conceptual frameworks (e.g., Corenblit et al. 2008; Corenblit and Steiger 2009; Gibling and Davies 2012). I will address and rectify this omission in this paper.

**Figure 1.**
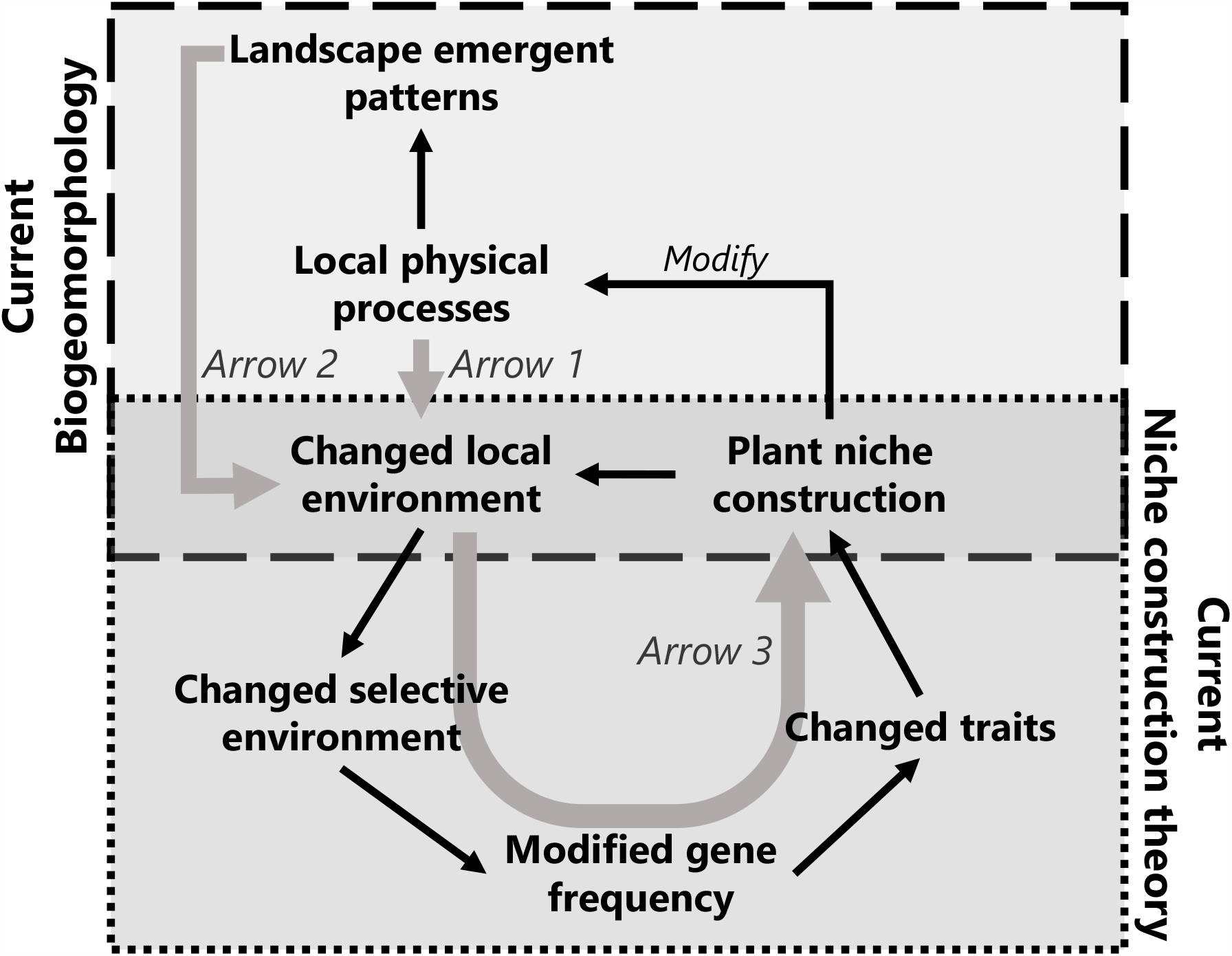
Integrating niche construction theory and biogeomorphology. Biogeomorphology studies plant niche construction interacting with physical processes at the local scale to give rise to patterns at the landscape scale. It assumes niche construction traits to be invariant in time. Niche construction theory concerns the effect of environment-altering activities that feed back on the evolutionary dynamics of plants. It often assumes that the environment only changes by organismal niche construction or assumes simple linear or periodic environmental changes. Integrating adjustments by physical processes at the local scale to niche construction (*Arrow 1*) and feedbacks from emergent properties at the landscape scale to niche construction traits (*Arrow 2*) is novel to niche construction theory. Integrating evolutionary feedback (*Arrow 3*) is novel to biogeomorphology.

Many biogeomorphic landscapes are self-organized; that is, complex patterns at large spatial scales can arise from mere local interactions (Murray, Goldstein, and Coco 2014). As such, effects of niche construction activities can propagate beyond the local scale at which they operate, via their interaction with local morphodynamics, and lead to emergent properties at the landscape scale (Fig. 1). For example, Schwarz et al. (2018) have shown that colonization rate, a life-history trait of plants in tidal marshes, not only influences sediment trapping locally, but also determines the ultimate steady state of the larger system. Colonization by fast colonizers favors stabilization of pre-existing channels. In contrast, dominance by slow colonizers facilitates the formation of new channels more broadly and thereby actively facilitates landscape self-organization. This is a typical occurrence in biogeomorphology, where niche construction interacts locally with physical processes, leading to emergent patterns globally (Fig. 1). While niche construction increases environmental predictability for organisms, the emergence of patterns beyond the local scale would potentially generate a set of entirely different selection pressures, feeding back to foster distinct evolutionary responses by niche constructors (Matthews et al. 2014). The implication of such cross-scale feedbacks to evolutionary dynamics of niche construction have not been examined heretofore (Fig. 1).

The role of independent environmental changes (changes independent of those brought about by niche constructors, e.g., physical processes such as sediment deposition, weathering, and erosion during landscape development) on evolutionary dynamics of niche construction, is not often acknowledged by niche construction theory (Laland, Odlingsmee, and Feldman 1996). The theory often assumes that all environmental changes are brought about by organismal niche construction (Fig. 1). The niche construction models that do consider independent changes, assume simple, linear changes (e.g., constant resource depletion or resource renewal rate), without considering the continual environmental adjustments that would occur in response to niche constructing activities. Furthermore, the models assume that the outcome of niche construction is homogeneously distributed to every organism in the population (Laland, Odling-Smee, and Feldman 1999; Silver and Di Paolo 2006). Current niche construction theory based on these simplifications suggests that across a broad range of conditions, niche-construction traits will drive themselves to fixation by hitchhiking on other traits favored in constructed environments (Odling-Smee et al. 2003). An initially rare recipient trait whose selection depends on niche construction could establish in otherwise unfavorable environments by forming spatial clusters of niche constructors. This creates environmentally mediated genotypic associations between the recipient trait and niche construction trait (Odling-Smee et al., 2003; Silver and Di Paolo 2006). During the development of biogeomorphic patterns, complex, nonlinear morphodynamics prevail and large spatial scale emergent properties arise from localized interactions. Morphological development encompasses rich, complex facets that are potentially of evolutionary significance: e.g., the time scale of landscape development (*vs*. plant lifespan), its style of change (Gibling and Davies 2012), spatial characteristics of topographic features (*vs*. species dispersal modes), the natural course of morphological development under physical constraints, and emergent patterns at the landscape scale. Current niche construction theory does not deal with this complexity. It remains to be determined whether plant niche construction traits can evolve in complex biogeomorphic landscapes, if these more realistic, complex environmental changes (here, landscape development) are accounted for.

In this study, I construct a geo-evolutionary model, which couples a population genetic module with a landscape evolution module, to generate multiple replicate trajectories of evolutionary and landscape change over several thousand-year periods. By manipulating (or eliminating) selected processes described above or feedbacks illustrated in Figure 1 in simulations, I can determine the consequences of failing to thoroughly integrate all elements of biogeomorphology and niche construction theory into a unified whole. In particular, I will use this approach to investigate two broad questions: (Q1) How does the evolution of niche construction traits influence landscape form and development? Does concurrent evolution of plant niche construction traits change the landscape steady-state outcome? If they occur, do resultant differences involve small changes in landscape elevation, or are difference drastic and involve landscape spatial pattern (structural difference). These questions are addressed by comparing the model results with and without concurrent evolutionary dynamics (Fig. 1–*Arrow 3*). In reciprocal fashion, (Q2) how does landscape development influence the evolution of niche construction traits? I examine (*a*) whether plant niche construction traits can still evolve when embedded in developing complex landscapes. If not, why not? If yes, (*b*) how does landscape development modify the evolutionary trajectory of niche construction traits? These questions will be addressed by comparing model outcomes with and without inclusion of landscape development and feedbacks (Fig. 1–*Arrow 1* and *2*).

## 2. Methods

### 2.1. The Model

#### Overview of the model

I base my analysis on a lattice-structured model, which couples landscape development with evolution of population genetics to analyze the joint evolution of a niche-constructing trait *G*, and a recipient trait *R*. Such designation of traits is common in the niche construction literature (e.g., Laland, Odling-Smee, and Feldman 1999; Silver and Di Paolo 2006). Greater *G* corresponds to stronger sediment trapping capacity by *G*, defined by the increase in sediment elevation per unit of plant biomass per unit time. *R* determines the elevation preferred by plants. Each plant has both traits—it can modify the environment by niche construction, and it also has genes to specify its own optimal environmental conditions. Since the environment is constantly changing, seldom do the environmental values specified by *R* and actual conditions coincide. This discrepancy drives selection and evolution. *G* and *R* take on continuous, positive values (quantitative traits). Changes in elevation are determined by not only plant niche construction activities, but also by abiotic processes, such as erosion, sediment deposition and diffusion.

#### Generality of the model

Model components—both landscape development and population genetics—are simplified to explore general, as opposed to system-specific elements of biogeomorphology and niche construction theory. In forming landscapes, some abiotic processes are not modifiable by any particular organism over a time scale necessary to effect a change in selection pressures or to generate an evolutionary response in a recipient organism (Matthews et al. 2014). Examples include rise of mountain ranges by tectonic activities or volcanic eruptions. Other abiotic processes such as weathering, erosion, and sediment transport are subject to effects of organismal, especially vegetative, activities. The model presented here focuses on those processes occurring in the critical zone—the layer of the Earth surface where abiotic and biotic processes actively interact (Brantley, Goldhaber, and Vala Ragnarsdottir 2007). Local elevation is the primary dependent variable that can be used to describe change in vast areas such as forests, aridlands, wetlands, and tundra etc. Elevation here is used as a surrogate for resource levels or other environmental conditions upon which plant growth relies. For example, sediment accretion reduces inundation stress for *Spartina anglica*, and enhances soil drainage and nutrient availability (Van Hulzen, Van Soelen, and Bouma 2007). For some beach grass species, growth is stimulated by sand burial in response to changes in soil temperature, space for root development, and increased nutrient and moisture availability (Martínez and Moreno-Casasola 1996; Maun and Perumal 1999). Diatom aggregations stabilize and accumulate sediments, forming elevated hummocks, creating an environment resistant to erosion (Weerman et al. 2010). In all these examples, changes in elevation are correlated with changes in attributes of ecological significance (e.g., soil chemistry, erosion, temperature, inundation) that will in turn influence plant survival and production. Other models that explicitly convert elevation to more-relevant ecological variables are possible, but broad features of biogeomorphic change will remain the same. Finally, and importantly, elevation reflects and determines the topographic contour of the Earth’s surface, which is the key descriptor of geomorphic form.

#### Initial condition

The model includes five state variables, plant age (*Age*), biomass of each plant (*Biom*), value of the geomorphic trait (*G*), value of the recipient trait (*R*), and elevation (*h*). Consider a landscape as a lattice-structured habitat consisting of an infinite number of sites. Each site can be in one of two possible states: occupied by a plant (vegetated) or unoccupied (bare). In the initial condition, I arbitrarily assign 70% of sites to be occupied by plants of ages that are also randomly sampled from the range of 1 and *Age*_*max*_. Each plant is further assigned a biomass value, as a function of its age. Biomass at site *i* in the initial condition is expressed as:

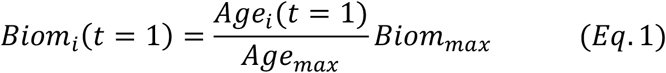

Each plant is assigned another two attributes: *G* and *R*. I assume that traits take on continuous, positive values within a range determined by the corresponding underlying alleles. *G* is sampled from a log-normal distribution with parameters *G*_*0*_ and σ_*G*_. Similarly, *R* for each plant is sampled from a log-normal distribution with mean *R*_*0*_ and variance σ_*R*_. The choice of distributions (e.g., normal distribution) does not qualitatively change the results reported in this study. Definition of parameters and their values used in the reference model are provided in Table S1. The initial elevation, *h*, is zero everywhere.

#### Dynamics of plants

Each plant grows (increase in biomass at the site it occupies), produces seeds annually after reaching the sexually mature stage, and eventually dies with an increasing mortality rate as the plant approaches its maximum age. The annual growth rate of a plant is influenced by the plant’s fitness to the local environment and the cost of niche construction. Plant growth rate is described as:

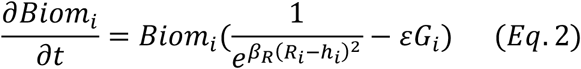

where 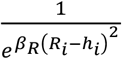 describes the effect of local environmental condition on plant growth, which is calculated as a function of the difference between the elevation preferred by plants specified by *R* and the actual elevation at site *i*. The closer these two values are, the higher the growth rate. *β*_*R*_ is a constant to regulate the rate of change in growth rate responding to the mismatch between the actual and the preferred elevation. The second term *εG* describes the cost for plants to invest in niche construction traits. *ε* is a scaler parameter. Plant age increases by one year with each time step. As a plant approaches its maximum age, mortality rate increases:

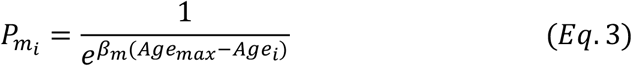

where *β*_*m*_ is a constant to moderate the rate of change in the mortality probability as a function of age. I use *Age*_*max*_ = 30 years in the reference model (Table S1). To examine the effect of plant lifespan, another two levels are explored, i.e., 400 and 1500 years (e.g., considering long-living trees modifying landscape topography by accelerating bedrock weathering via root respiration, e.g., (Dong et al. 2018)).

#### Seed production and dispersal

After a prescribed age, *Age*_*s*_, plants begin to produce seeds. Seeds are assumed to be produced annually. The number of seeds produced annually is proportional to the biomass of the parent plant in that year. Seeds then disperse from the source plant. The probability that a seed from a given plant (site) lands on an unoccupied site and successfully germinates is determined by (1) the Euclidean distance between the source and the destination site, (2) the biomass of the parent plant, which determines the number of seeds produced by that plant, and (3) the suitability of environmental conditions at the landing site for the seed. The suitability is measured by the difference between *R* of the seed and elevation of the landing site, *h*. Genotypes of the seed are assumed to be the same as the parent plant (the model assumes perfect heritability, i.e., the phenotype is exactly the genotype with environmental variance or plasticity), unless mutation occurs (described below). When the value of *R* and *h* are close, it means the elevation of the landing site is close to the optimal elevation required for seed germination. To sum up, the probability for a plant at site *i* to have its seed successfully disperse to an unoccupied site *j* and germinate, *p*_*ij*_, is described by:

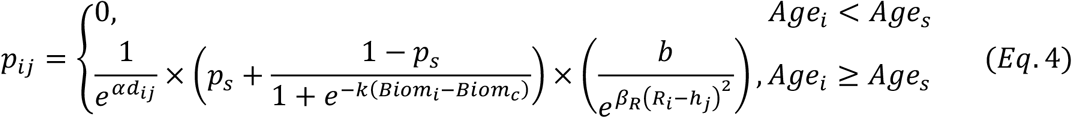

where *d*_*ij*_ is the Euclidean distance between source site *i* and destination site *j. p*_*s*_ is the minimum probability for a plant older than age *Age*_*s*_ to produce seeds that year (*p*_*s*_ is positive value, assuming that once the plant reaches its sexual maturity, it will produce every year). *Biom*_*c*_ is the critical biomass above which the increase of seed production as a function of biomass slows down. *α* is a shape parameter characterizing dispersal mode. Three levels of dispersal are investigated, i.e., *α* at 0.7 (reference model), 0.1, and 0.001. These levels correspond to taking 3, 30, 3000 cells’ distance for the likelihood of seed dispersal to drop to 5% (the system size is 100 ⨯ 100 cells), ranging from very local to very long-distance dispersal. *b* is a constant to regulate the rate of change.

#### Mutations

With a small probability, mutations can independently occur to *G* and *R*, when seeds are produced. When a mutation occurs, the trait value of the seed will be different from the trait value of its source plant. The mutant *G* or *R* is sampled from the same lognormal distributions of *G* or *R* applied in the initial condition (see ‘*Initial condition*’, above).

#### Landscape development

Elevation change is jointly determined by abiotic processes and plant niche construction. I assume a baseline rate of elevational change caused by abiotic processes globally, *J* (e.g., baseline sediment deposition rate). The effect of niche construction on elevation change is expressed as a function of biomass and *G*. A hump-shaped relationship between rate of elevational change and elevation is assumed (Strudley, Murray, and Haff 2006). The elevation change is slow at the beginning, accelerates, and as the landscape approaches a steady state asymptotically, becomes slow again. Furthermore, to describe the morphological consequence beyond the local scale, I assume that the accumulation of sediments in one place induces negative effects on sedimentation in sites nearby, *S*. Such long-distance negative feedbacks are common in ecosystems (Rietkerk and van de Koppel 2008; Murray, Goldstein, and Coco 2014). For example, turbulence and enhanced erosion lead to deep gullies around elevated vegetation patches in fluvial environments, which reduces the likelihood for other plants to establish; that is, a flow divergence effect occurs (Cornacchia et al. 2019; Zhao et al. 2019). Long-distance negative feedbacks are essential to form regular patterns in many ecosystems (Rietkerk and van de Koppel 2008). I refer to *S* as the force of pattern formation hereafter. Lastly, the model includes sediment diffusion. The change of elevation is described as:

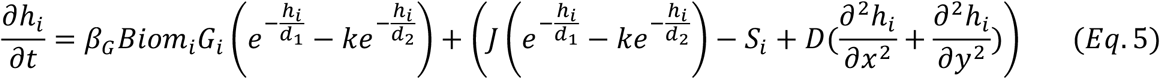

where *D* is the diffusion coefficient of sediments and *d*_*1*_ and *d*_*2*_ are the decay scaling factors. *k* is a dimensionless coefficient (*k* < 1) that determines the magnitude of the rate when *h* = 0, relative to (*J* ± *β*_*G*_*Biom*_*i*_*G*_*i*_) (Strudley, Murray, and Haff 2006). *S* is the long-distance negative suppression caused by the existence of local high patches on the increase of elevation nearby. Each site in the system receives negative effects from all the sites that are higher than the focal site, the effect size being negatively correlated to the Euclidean distance between the higher site and the focal site. The negative effect received by site *i* is described by:

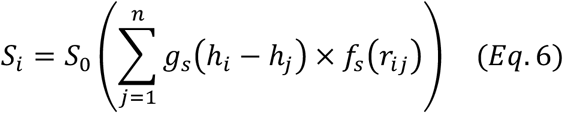

where the summation is over the entire system and *n* is the total number of sites in the system. *S*_*o*_ is the negative effect a site received from another site which is one-unit higher and one-unit distance away. The function *f*_*s*_ defines the amount of influence a higher site that is *r*_*ij*_ away has on a lower site. A Gaussian function is assumed for *f*_*s*_*(r)* with a rate coefficient *ω*:

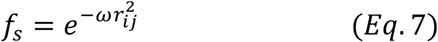

*g*_*s*_ in *Eq*. 6 is defined as zero, when site *j* is lower than site *i* and *g*_*s*_ is equal to the difference in elevation otherwise:

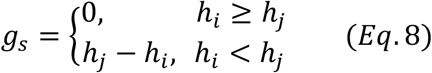

### 2.2. Numerical Experiments

The models were run on lattices of 100 ⨯100 cells with periodic boundary conditions (probabilistic cellular automaton), with each cell representing a site. The time step in the model is one year and I ran the model for 100,000 years to ensure that the landscape morphology and evolutionary dynamics reach a statistically steady state. The model is solved numerically using an implicit finite difference method. The predictions on the effect of geo-evolutionary coupling on landscape morphology and on evolution of niche construction are tested with numerical experiments.

#### Effect of evolution on geomorphology

To investigate the effect of evolution on landscape morphology, I created a gradient of geo-evolutionary coupling. The strength of coupling is manipulated by the constants multiplying *G* (*β*_*G*_ in *Eq*. 5) and *R* (*β*_*R*_ in *Eqs*. 2 and 4). Three levels of *β*_*G*_ are 0.00001, 8, and 30 (reference model). Small *β*_*G*_ describes a system where landscape morphology is dominated by abiotic processes. This occurs when abiotic processes greatly outpace plant-induced elevational changes or when abiotic processes are not modifiable by organisms. Increasing *β*_*G*_ reflects an increasingly important role of plant niche construction in landscape change. The sensitivity of plants to elevation is represented by the decay scaling factor *β*_*R*_ in *Eqs*. 2 and 4. Three levels of sensitivity are constructed, with *β*_*R*_ being 0.4, 2, and 10 (reference model). A small *β*_*R*_ means plants are insensitive to elevation, in which case even a large difference between *R* and *h* does not have much effect on plant growth (*Eq*. 2) or on the success rate of seed germination (*Eq*. 4). Geo-evolutionary coupling is strongest when plants have a strong effect on changing landscape development and the survival and reproduction of plants are highly sensitive to resultant elevational changes. Simulations with these 3 ⨯ 3 parameter combinations allow me to analyze the effects of the different degrees of geo-evolutionary coupling.

Furthermore, to explore potential nonlinear changes in landforms, I performed a systematic analysis of a wider parameter domain. I created a strong pattern formation scenario (*S*_*0*_ = 0.012) and a weak one (*S*_*0*_ = 0.0032; used in the reference model). Under strong pattern formation, I created 15 levels of *β*_*R*_ between 0 and 12 and 15 levels of *β*_*G*_ between 0 and 32 (225 combinations). For the scenario of weak pattern formation, I created 19 levels of *β*_*R*_ between 0 and 15 and 19 levels of *β*_*G*_ between 0 and 40 (361 combinations). In particular, when *β*_*G*_ = 0, plants niche construction activity is zero. When *β*_*R*_ = 0, plant growth or reproduction is not influenced by elevation at all, which represents the case of no selection or evolution. The effect of including evolution on landscape development therefore can be assessed by comparing the results of models where *β*_*R*_ ≠ 0 with the models where *β*_*R*_ = 0.

#### Effect of landscape development on evolution

To investigate the effect of simultaneous landscape development on the evolution of niche construction, I created model scenarios by turning on and off the various combinations of physical processes involved in landscape development. Scenarios are as follows: (1) The reference model includes all the physical processes—baseline elevation change, long-distance negative feedback, and sediment diffusion; (2) all physical processes are turned off; (3) only sediment diffusion is turned on; (4) only baseline elevation changes are turned on; (5) both diffusion and baseline elevation change are turned on; and (6) diffusion and long-distance negative feedback are turned on. Effects of different values of the diffusion coefficient *D* are also considered. These manipulations allow me to not only assess the effect of simultaneously occurring landscape development on the evolution of niche construction, but also to identify the evolutionary consequences of specific physical processes.

For each numerical experiment, I use 10 or 20 replicates to capture the mean, standard deviation, and statistical significance of results. With each simulation outcome, I analyze the frequency distribution of trait values, population-level mean trait values, correlation between *G* and *R* at the landscape scale over time, and the evolution rate of *G*. Evolution rate is quantified in two ways: (1) change of the population mean *G* divided by time (*v*_*T*_, with unit *G* yr^-1^); and (2) change of the population mean *G* divided by the change of the population mean *R* in the same period (*v*_*R*_, with unit *G R*^-1^). Further, I quantify mean population fitness, approximated by the landscape-scale mean probability of successful seed establishment (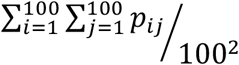; *Eq*. 4), which integrates the number of offspring produced and the fitness of seeds to the environment. Lastly, I use semivariance to characterize the patch size and Moran’s I for spatial autocorrelation to compare the spatial patterns of *R* and *G*.

## 3. Results

### 3.1. Landscape morphology

Allowing plants to evolve always changes the resulting landscape form and development. The change could be a small difference in steady-state landscape elevation or could be drastic differences in landforms (Fig. 2), but evolution always matters. When the force of pattern formation is relatively weak, the landscape evolves to a low-relief (slope in 10^−4^) and high-elevation state (Fig. 2E). In this state, the steady-state landscape generated by the model with plant evolution increases the steady-state landscape elevation and reduces its relief, in comparison to the without evolution case. With increasingly stronger geo-evolutionary coupling, the steady-state landscape shifts toward even higher elevations and lower relief (Fig. 2E). As the force of pattern formation strengthens, the landscape evolves to either the regularly patterned state if plants have low niche construction capacity and high sensitivity to elevation or to the high-relief state (slope in 10^−3^) if plants have high niche construction capacity and low sensitivity to elevation (Fig. 2). By including evolution of plant niche construction traits in the model, the likelihood of forming regularly patterned landscapes is enhanced (Fig. 2D). Allowing plants to evolve reduces the steady-state landscape elevation and its relief. (Fig. 2D2). With increasing sensitivity of plants to elevational changes, and/or decreasing niche construction activities, the landscape shifts from a high-relief landscape to a regularly patterned landscape (Fig. 2). Diverse outcomes are possible.

**Figure 2.**
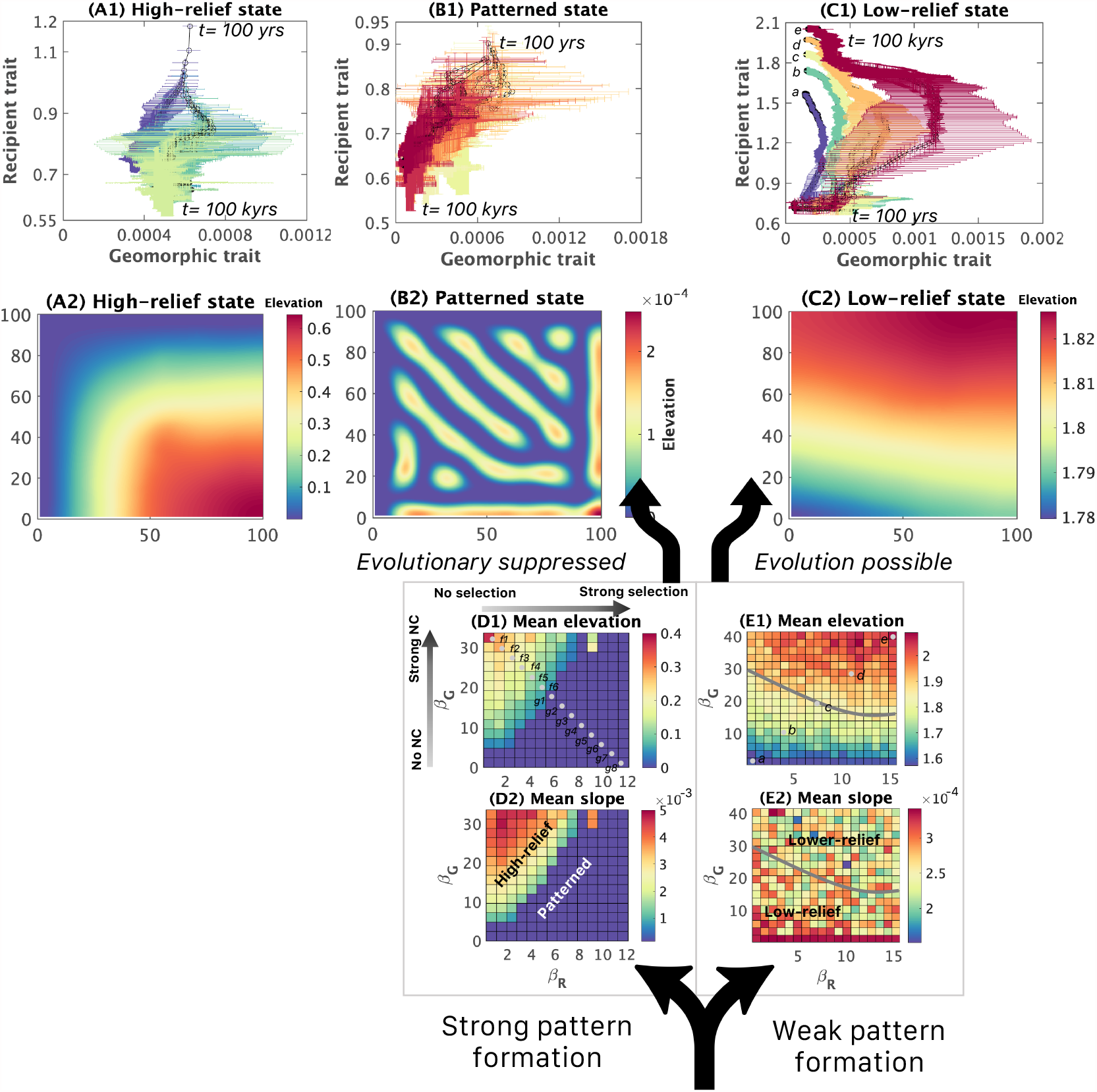
Phase transition of landscape morphology influenced by coupled landscape development and evolution of niche construction. When the long-distance negative effect (force of pattern formation) is relatively weak (*S*_*0*_ = 0.0032), the landscape develops into a low-relief state (C2). With increasingly strong geo-evolutionary feedback, the steady state of the landscape features a higher elevation (E1) and a lower relief (E2), and the niche construction trait can evolve (C1). When long-distance negative feedback is relatively strong (*S*_*0*_ = 0.012), the landscape develops into high relief landscape (A2) or patterned state (B2), depending on the strength of geo-evolutionary feedback. These two states are evolutionary suppressors of niche construction traits (A1 and B1). (D1) and (E1) show the steady-state mean elevation, and (D2) and (E2) are the steady-state mean slope. (A1), (B1), and (C1) depict the change of the population mean trait value over 100,000 yrs. (A1) shows trait evolution at six different levels of geo-evolutionary feedback strengths (represented by *f*1-*f*6 in (D1)). (B1) shows trait evolution at eight levels of geo-evolutionary feedback strengths (*g*1-*g*8 in (D1)). Similarly, (C1) shows trait evolution at five levels of geo-evolutionary feedback strengths (*a, b*, …, *e* in (E1)). At each strength level, the mean and standard deviation of population mean trait value were constructed using 10 replicated numerical simulations.

### 3.2. Alternative evolutionary trajectories

*R* always tracks the elevational change (Fig. 3A), while evolutionary dynamics of *G* are sensitive to landscape development (Fig. 2). Selection on *G* is effective before the landscape reaches its steady state; however, whether *G* can continue to evolve after the landscape reaches its steady state depends on the landscape spatial configuration (Fig. 2). If the landscape stabilizes at the low-relief state, selection on *G* continues to operate (Fig. 4D3). In the early stage of landscape development to this state, *h* is much lower than the level preferred by plants (Fig. 3A). Niche construction reduces the difference between population mean *R* and *h*, benefiting plant growth. This is positive niche construction (defined as niche constructing acts that, on average, increases the fitness of the niche-constructing organisms; Odling-Smee et al. 2003). Plants that have higher niche construction capacity will be selected for. As expected, model results show that *G* increases (after the initial decrease in *G* caused by the initial condition effect) (Fig. 3). The diversity of *R* is low in this phase (Fig. 3E), because the elevation is near the lower limit of the species’ tolerance and the difference between actual and preferred elevation is large, indicating high selection pressure. Hence, any mutation of small value *R*, once it appears, will rapidly spread across the landscape. It takes ∼180 generations for *R* to evolve close to *h* and *h* starts to exceed *R* in some places of the landscape (Fig. 3). The system enters the local positive niche construction phase. Diversity and population mean value of *R* increase in this phase (Fig. 3D–E). Large value *R* (*R* > *h*) and large value *G* form niche constructing spatial clusters. As a result, population mean *G* continues to increase. With further landscape development, *h* begins to approach the upper limit of the species’ tolerance, and mutations of *R* large enough to match *h* become rare. The proportion of the landscape where *h* is greater than the elevation preferred by plants becomes dominant—that is, the system enters the negative niche construction phase (refers to niche-constructing acts that, on average, decreases the fitness of the niche-constructing organisms; Odling-Smee et al. 2003). In this phase, plants benefit from lower niche construction activity, since *h* is already higher than the elevation preferred by plants. Hence, plants of large value *R* and small value *G* will be favored. As a result, a trend of increasing population mean *R* and decreasing *G* is observed (Fig. 3D). Meanwhile, diversity of *R* in the population decreases (Fig. 3E). At the very late stage of landscape development, the population is dominated by only a few (< 3) *R* (Fig. 3E). However, the diversity of *G* is much higher than *R* and stays at about the same level over time.

**Figure 3.**
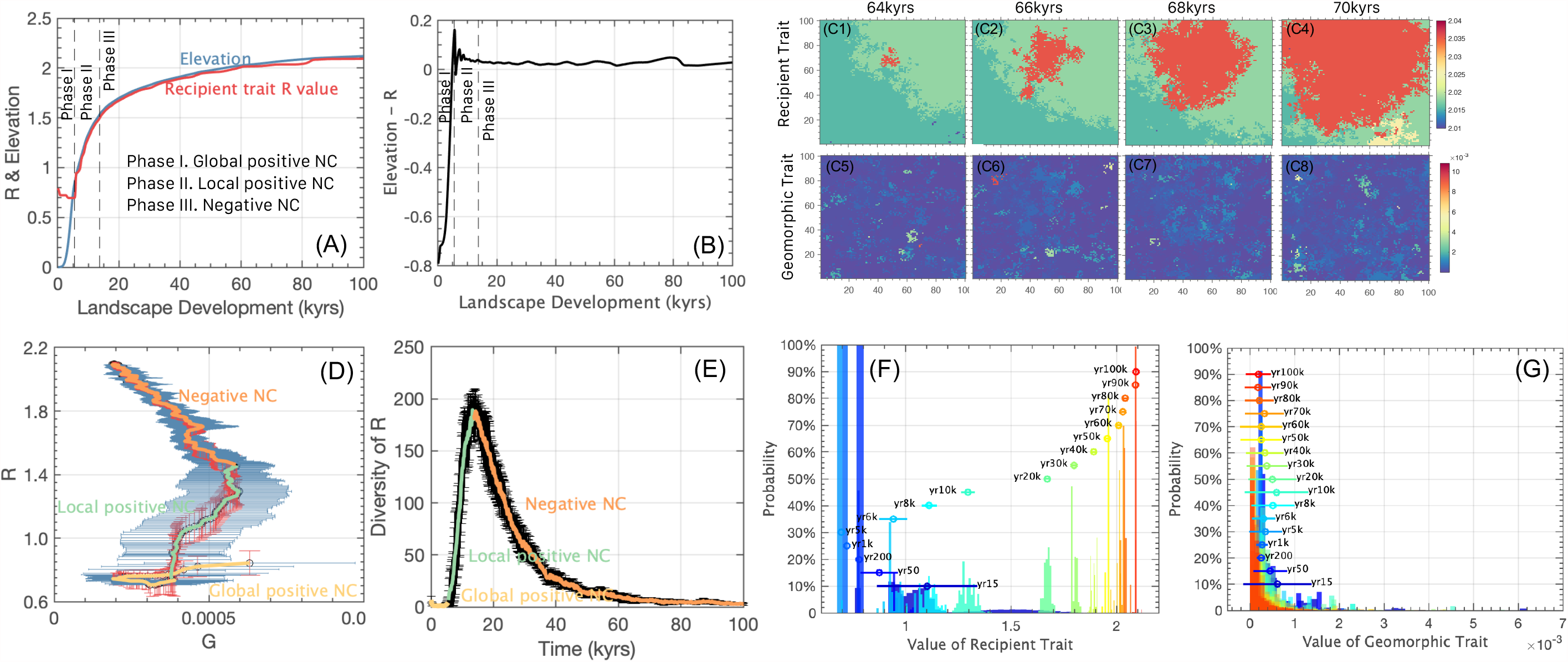
Evolution of the recipient trait *R* and the geomorphic trait *G* over 100 kyrs landscape development. (A) and (B) describe the change of mean elevation and population mean *R* over 100-kyr simulation time. This period can be divided into three phases: global positive niche construction, local positive niche construction, and negative niche construction. C1-C4 shows the formation and expansion of a recipient trait cluster. Such clustering is not observed in the pattern of *G* (C5-C8). (D) describes the evolution of *G* and *R* with standard deviation constructed by 10 replicated simulations, labeled with the three phases. Transition from local positive niche construction phase to negative niche construction phase occurs when diversity of *R* in the system starts to decline, as described in (E). (F) and (G) are histograms showing the evolution of *R* and *G* over time, labeled with the landscape-level mean and standard deviation of trait values for selected years.

**Figure 4.**
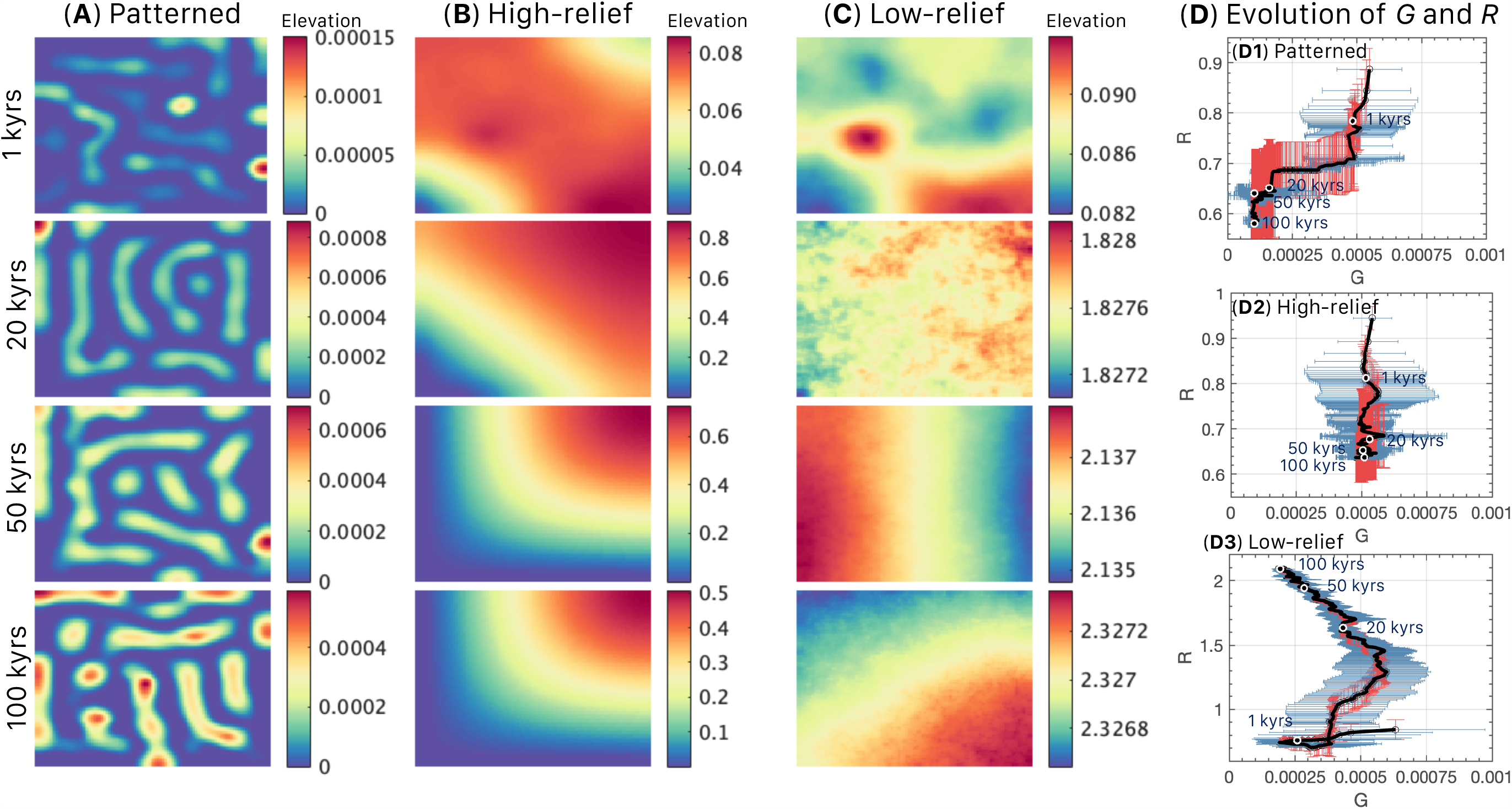
Three alternative stable states of landscape morphology and their development over time (A–C). (D1)–(D3) show the evolution of the population mean trait value for geomorphic trait *G* and for recipient trait *R* over time in the three alternative landforms. The low-relief landscape (C) is generated by the reference model (*S*_*0*_ = 0.0032, *β*_*R*_ = 10, *β*_*G*_ = 30, see Table S1 for values of the other parameters), patterned landscape (A) is generated by relatively strong long-distance negative feedback (*S*_*0*_ = 0.012), with *β*_*R*_ = 5.3, *β*_*G*_ = 4.8, and the high-relief landscape (B) is also generated by relatively strong long-distance negative feedback (*S*_*0*_ = 0.012), but with *β*_*R*_ = 2.75, *β*_*m*_ = 21.6.

The correlation between *G* and *R* in these phases is low overall (mean of 0.06, with a standard deviation of 0.11 over the course of landscape development) (Fig. S1). The correlation is lowest during the first phase—global positive niche construction, limited by the low value *R* mutants (Fig. S2). Correlation increases during local positive niche construction. As *h* continues to increase, mutation of *R* that can match *h* rarely occur, since *h* approaches the species’ upper limit. As a result, with the system entering the negative niche construction phase, diversity of *R* in the population begins to decline, and the association between *G* and *R* breaks down (Fig. S2). Geo-evolutionary feedback enhances the correlation between *G* and *R* (mean correlation is 0.06 *vs*. 0.00, under strong and weak geo-evolutionary coupling respectively) (Fig. S1).

For landscapes that develop to high-relief or regularly patterned landforms, before the steady state is reached, the evolutionary dynamics of *G* are the same as described above for a low-relief landscape. Once the steady state is reached however, selection on *G* stops (Fig. 2A1). In a high-relief state, sediment trapping by *G* is offset by fast sediment diffusion caused by high relief. As a result, even though large value *G* is favored, it is not possible to increase its frequency in the population. In a regularly patterned steady state, selection on *G* ceases to be effective, even though mutations of large value *G* are still favored, since *h* is much lower than the elevation preferred by plants (Fig. 4). The inability of *G* to evolve is caused by the particular landscape configuration. When a mutation of large value *G* occurs on the ridge of the landscape, diffusion offsets the effect of sediment trapping by plants. If mutations of large value *G* occur in valleys, sediment accumulation is suppressed by the long-distance negative feedback exerted by relatively high ridges nearby. As a result, both high-relief and patterned landscapes are suppressors of natural selection on *G*.

### 3.3. Stochasticity in evolutionary outcomes

Regardless of the distinct evolutionary trajectory caused by alternative landforms, stochasticity in the evolution of *G* is high and is overall much higher than that in the evolution of *R* (Fig. 3D and G). Long lifespan and long-distance dispersal of plants greatly amplifies stochasticity to such an extent that it overwhelms selection on *G* (Fig. 5). There is no general trend of increasing population mean *G* in the positive niche construction phase, nor is there a descending trend in the negative niche construction phase (Fig. 5B–C). This is true even when the landscape development is turned off (Fig. S3). The evolution of *R* however is not influenced, and *R* always tracks the environment (Fig. 5). When dispersal is local, seeds are more likely to land on nearby sites of similar *h*. In this case, the number of seeds produced, which reflects fitness of parent plants (*Eq*. 4), has a strong effect on which seeds get established in vacant sites. In contrast, with long-distance dispersal, the condition (elevation) of the landing site plays a stronger role in determining success of seed establishment. Hence, stochasticity in the evolution of *G* will be much higher (Fig. 5). Traits of short-lived plants are updated more frequently and can therefore more closely track the changing landforms, while longer lifespan amplifies the role of stochasticity (Fig. 5).

**Figure 5.**
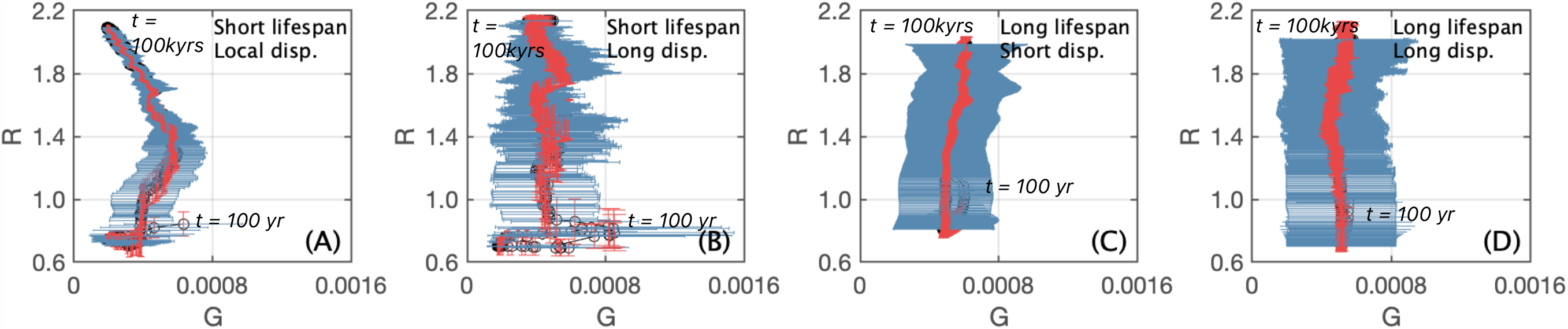
Effect of dispersal mode and plant lifespan on the evolution of niche construction trait *G* and recipient trait *R*. Niche construction trait can evolve when dispersal is local, and plants’ lifespan is short. The black open points represent the landscape-scale mean trait value averaging over 10 replicated simulations. The standard deviation for *G* and *R* is calculated from 10 replicates. (A) shows the results of the reference model (parameter values provided in Table S1), with lifespan = 30 yrs, and local dispersal *(α*= 0.7). *α* = 0.01 is used for long-distance dispersal in (B) and (D), and lifespan of 1,500 years was used for long lifespan scenario in (C) and (D).

Physical processes in landscape development have process-specific effects on the evolution of traits. When baseline elevational change is turned off, stochasticity in the evolution of *R* increases (Fig. 6E–G). This is because baseline elevational increases impose a directional selection on *R*. Without the baseline rate, elevation can only increase by niche construction activities. Accelerated evolution of *R* increases the likelihood of forming large niche construction clusters, which reduces stochasticity in the evolution of *G* and increases its evolution rate (Figs. 6 and S4). Sediment diffusion reduces the evolution of *G* in both positive and negative niche construction phases (Fig. 6). This is because the larger elevation increase caused by larger *G* during positive niche construction and the reduced elevation increase by smaller *G* during the negative niche construction phase can both be offset by sediment diffusion. In both phases, the effect of the favorable *G* is dampened or even eliminated. Lastly, when all independent physical processes are turned off and the landscape is entirely controlled by niche constructing activities, the evolution rate of *G* in the positive niche construction phase is more than twice that in the reference model (13 × 10^−4^ *vs*. 6 × 10^−4^ *G R*^-1^; Fig. 6A and E), reflecting the effect of stochasticity introduced by landscape development on slowing down the evolution of *G* by preventing the formation of large niche construction clusters (Fig. S7).

**Figure 6.**
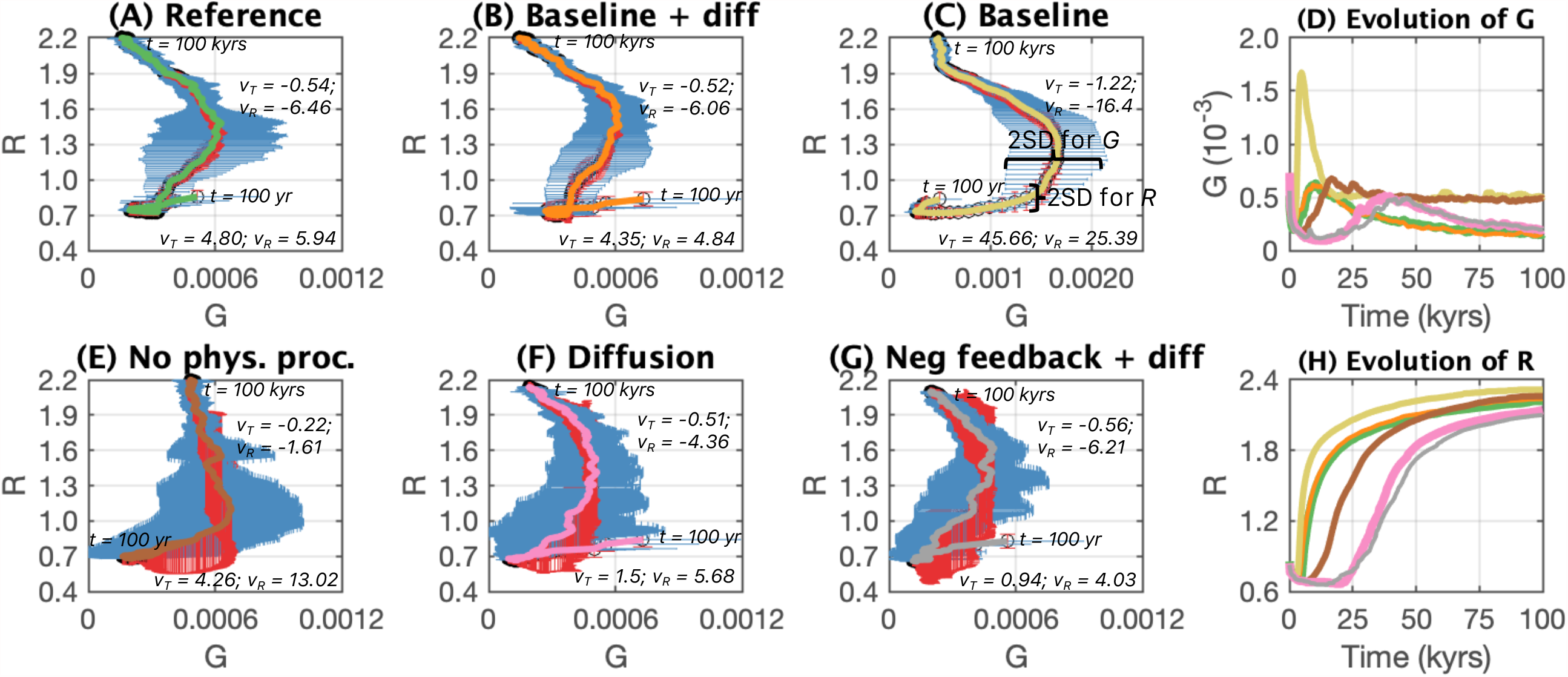
Effect of different physical processes on the evolution of niche construction trait *G* and recipient trait *R*. Results from (A) reference model with all the processes turned on; (B) when baseline elevation rate and sediment diffusion are turned on; (C) when only baseline elevational change is turned on; (E) when all the independent physical processes are off; (F) only diffusion is on; (G) only long-distance negative feedback and sediment diffusion are on. (D) and (H) describe the evolution of *G* and *R* over 100 kyrs under each of the six scenarios described. In (A)-(C) and (E)–(G), black open points connected by solid lines are the population mean trait values averaging over 20 replicated simulations, with standard deviations (vertically for *R* in red and horizontally for *G* in blue). *V*_*T*_ (10^−8^ *G* yr^-1^) and *V*_*R*_ (10^−4^ *G R*^-1^) are the phase-specific evolutionary rates of *G*, with positive values in the positive niche construction phase (*G* increases its values), and negative rates in the negative niche construction phase. In (D) and (H), the landscape mean trait value is averaged over 20 replicates, with color coding corresponding to the color of solid lines in (A)–(C) and (E)–(G).

### 3.4. Comparing geomorphic and recipient traits

*G* and *R* show very different patterns in evolution. For *R*, toward later stage of landscape development, a favorable *R* can persist much longer in the population (up to ∼750 generations) and become more spatially dominant (Fig. S5A). In contrast, for *G*, duration of a *G* and its spatial dominance do not change significantly over time (Fig. S5B), with a mean of ∼5 generations and mean maximum spatial coverage for a *G* of 2.1% (compared to 10.4% for *R*). Consequently, the genotype of *G* maintains a much higher diversity (∼50 unique *R* at any time on the landscape *vs*. ∼350 for *G*. The most dominant *G* reaches ∼38%, compared to 95% by a dominant *R*) (Fig. 3C).

The landscape-level mean correlation coefficient between *G* and *h* is highest, followed by that between *R* and *h* (Fig. S1). *G* and *R* have the lowest correlation, as the correlation between *G* and *R* is indirect, mediated by *h*. Patches of *G* and *R*, especially *R*, have sharp boundaries (Fig. S6). Results from semi-variance analysis indicate that the patch size of *R* is much larger than *G*. Moran’s I quantifies spatial autocorrelation. Its value is highest for elevation, followed by *R, G*, and lastly, biomass. Moran’s I for trait values are significantly positive, indicating spatial clustering. Moran’s I for *R* spikes in the early stage of landscape development when diversity of *R* is low (Fig. 7). Autocorrelation declines rapidly as diversity of *R* in the population increases (Fig. 7). Spatial autocorrelation of *R* reaches its lowest level when *R* diversity peaks, after which the autocorrelation of *R* gradually increases as its diversity decreases (Fig. 7). Geo-evolutionary feedback increases the spatial autocorrelation of both *G* and *R* as well as the correlation between the trait value of *G* and *R* (Fig. S1).

**Figure 7.**
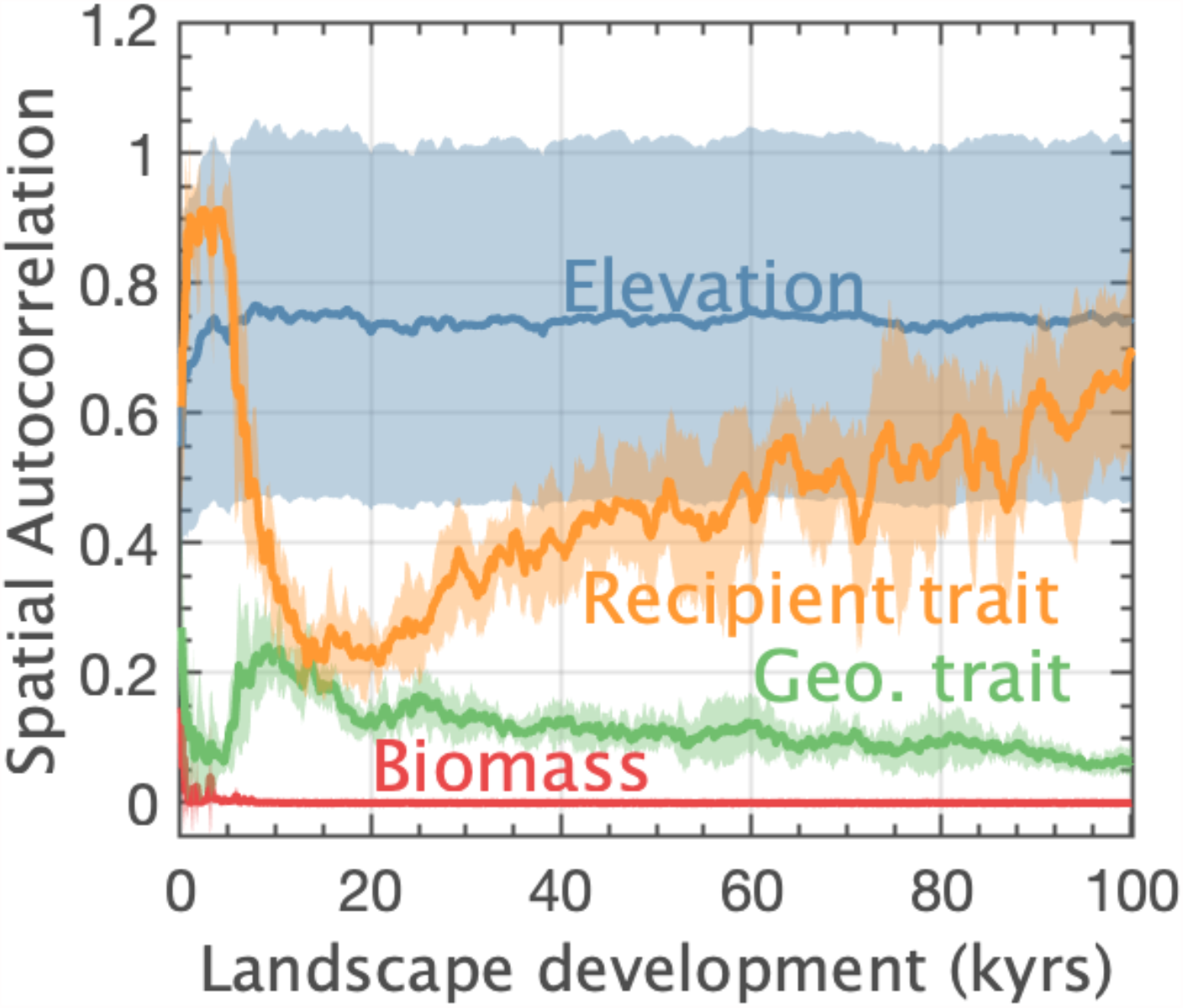
Mean spatial autocorrelation measured by Moran’s I. Landscape-scale Moran’s I of elevation, biomass, geomorphic trait, and recipient trait over the course of landscape morphological development. Confidence interval is constructed using 10 replicated simulations.

## 4. Discussion

This study demonstrates that coupling landscape development and evolution has significant implications for niche construction theory and for biogeomorphology (Fig. 1). Allowing plants to evolve during landscape development always results in different landforms compared to the case where evolution is ignored or excluded. The omission effect ranges from mild differences in steady-state elevations to drastic differences in landforms (Fig. 2). Allowing the landscape to change while evolution of niche construction occurs results in distinctively different evolutionary trajectories for niche construction traits. Some of these trajectories develop into states where natural selection is suppressed, and evolution stops. Below, I discuss the implications of geo-evolutionary coupling for (1) biogeomorphology, (2) niche construction theory, and (3) responses of biogeomorphic landscapes to large scale changes, such as in global climate.

### 4.1. Implications of evolving plants for biogeomorphology

Allowing plants to evolve in the model results in different landforms than if traits are considered to be static as has been done in past biogeomorphic studies. When the force of pattern formation is weak, the model scenario allowing plant evolution generates landscapes that stabilize at a state of higher elevation and lower relief (Fig. 2). The extra increase in elevation is a consequence of the evolution of niche construction traits that evolve toward higher values in the early, positive niche construction phase. Stronger geo-evolutionary coupling leads to landscapes that stabilize at even higher elevations (Fig. 2). When the force of pattern formation is strong, the system could stabilize at either patterned or non-patterned states without considering the evolution of plant traits (Fig. 2). Accounting for the evolution of plants however increases the likelihood that the system will stabilize at the regularly patterned state rather than the high-relief state. This is caused by combined effects of selection on *R* and the inability of *G* to evolve toward favorable values in the patterned state (Fig. 2). As a result, the total amount of sediment trapped by plants is much lower than if selection did not occur. This increases the likelihood for patterned landscapes to emerge. The likelihood of pattern formation is even higher as geo-evolutionary coupling strengthens.

While the topographic signature of life on Earth is unmistakable at almost all scales, Dietrich and Perron (2006) conjecture in their seminal work that the rise of life does not create new forms of topography on Earth. They do not consider simultaneous biological evolution. I find that allowing plants to evolve in the model also does not create new topographic signatures. Nevertheless, including evolutionary processes, which more realistically depicts system dynamics, always results in different landforms. The difference in landforms between cases with and without biological evolution can range from small differences in steady-state elevation, to drastic differences in the landform (e.g., from non-patterned to patterned landscapes) (Fig. 2).

The amount of difference is contingent on the form of landscape development (processes involved in landscape development) and the strength of geo-evolutionary coupling (Fig. 2). It is worth noting that the evolutionary dynamics considered here only involve changes in population genetics. It is reasonable to expect that innovation derived from evolution—such as mutations of radically novel traits—would create novel topography, especially in systems where evolution and landscape development are closely coupled and complementary information from biological evolution comes to dominate (Krakauer et al. 2020). Taking evolutionary dynamics into consideration renders states of future landscapes much less predictable.

### 4.2. Implications of landscape development for niche construction theory

#### 4.2.1. Contingency of evolutionary trajectories on landscape development

Niche construction often occurs at the local scale as organisms modify their immediate environment. As such, niche construction is perceived to make the environment more predictable for organisms (Laland, Odling-Smee, and Endler 2017). This is only supported when all of the independent physical processes of the model are turned off and landscape change is solely shaped by niche constructing activities. In that scenario, large niche constructing clusters form and the landscape develops into mosaics of patches of different elevations with abrupt boundaries (elevation difference up to ∼0.8 m; Fig. S7). In fact, niche construction can modify the local environment to the extent that elevation difference between patches is so large that environmental isolation arises. As a result, effective dispersal, hence gene flow, among patches is limited, leading to subpopulations of distinct *G* and *R* genotypic signatures (Fig. S7).

Once landscape development is turned on, the evolution of niche construction traits becomes much less predictable (Fig. 1). Multiple stable states in landscape formation are common, especially when the landscape is self-organized, in which case local interactions give rise to patterns at much larger spatial scales (Murray, Goldstein, and Coco 2014). Alternative stable landforms imply distinctly different evolutionary trajectories. Evolutionary graph theory, which generalizes population structure by arranging individuals on a graph, suggests that spatial structure influences the fixation rate of advantageous mutants (Lieberman, Hauert, and Nowak 2005). Some graphs act as suppressors and others as amplifiers of natural selection (Lieberman, Hauert, and Nowak 2005). Model results suggest that the coupled landscape-plant population system can develop into evolutionary suppressor states by themselves. In particular, when the force of pattern formation is strong, evolution of *G* occur until the steady state of high-relief state or regularly patterned state is reached. When that happens, selection ceases to be effective. In both states, natural selection on *G* is suspended as a result of canceling the effect of niche construction occurring in particular landscape spatial configurations. Sediment accrual by niche construction could be offset by diffusion when slope is relatively steep (Fig. 4A–B), or it could be offset by the enhanced erosion caused by nearby elevated geomorphic features (Fig. 4A). While these two evolutionary suppressors are specific to the model construct in this study, landscape configurations of other forms that can suppress or amplify evolution are likely in other models and in real systems. Without considering simultaneous landscape development, such evolutionary dynamics would not be revealed by niche construction theory.

When the pattern formation force is relatively weak, landscapes stabilize at a state of low relief, wherein selection is possible (Fig. 2E). Even though selection does not operate directly on *G* in the model, niche construction theory predicts that *G* can drive itself to fixation through hitchhiking on other traits favored in constructed environments (Laland, Odling-Smee, and Feldman 1999; Lehmann 2008; Silver and Di Paolo 2006). However, I find that the statistical association between *G* and *R* is very low (Fig. S1), indicating the low effectiveness of hitchhiking (Fig. S7B). Further, evolution of *G* is sensitive to physical processes in landscape development, as demonstrated by numerical experiments of turning on and off different processes (Fig. S4). First, the long-distance negative effect, which is considered to be essential to form regular patterns in many ecosystems (Rietkerk and van de Koppel 2008), determines the possible domain of evolutionary dynamics of niche construction traits—whether the population will be trapped in an evolutionary suppressor state (Fig. 2). Diffusion of sediment, another abiotic process in the model, moves sediments from higher elevation sites to lower neighboring sites. If plant niche construction occurs and increases the elevation at a site, sediment diffusion distributes this outcome of niche construction to its lower neighbors. As a result of this “imposed altruism”, the elevation of neighboring sites can also increase, even without niche construction. It has been shown that plant niche construction evolves only when dispersal is limited (Kéfi et al. 2008). Here I show that if sediment diffusion is rapid, and the cost of *G* is high, favorable *G* will not increase its frequency, even under local dispersal (Fig. S8). The third abiotic process in the model, baseline elevational change, accelerates the evolution of *G*. The rate of baseline change could reflect sediment supply. An increase in sediment supply has been shown to promote a shift from Brownian to Levy clonal expansion in dune building grasses, maximizing sand trapping efficiency (Reijers et al. 2020). I find that sediment supply could further influence the rate of evolution of niche construction traits. Physical processes in the model that describe landscape development here are by no means comprehensive. In natural systems, the complexity of morphodynamics is unusually high. However, results from this set of numerical experiments at least show that to understand evolution of plant niche construction, it is necessary to consider simultaneous environmental changes (here landscape development). Furthermore, specific physical processes could modify—even completely alter—the evolutionary trajectory of niche construction traits. Studies in niche construction theory so far, which assume independent environmental change to be nonexistent or simply linear or periodic, could have led to these distorted or worse, inaccurate, conclusions.

#### 4.2.2. Stochasticity overwhelms selection

Long lifespan of plants and/or long-distance dispersal drastically amplify stochasticity in the evolution of *G*, such that it overwhelms the effect of natural selection on niche construction traits (Figs. 5 and S3). Long-distance dispersal reduces the success of seed establishment, as the condition of the landing site can be vastly different from the source. It is worth noting that while treated as an invariant trait in my model, and in reality, dispersal is a trait that can evolve with landscape change. Plant dispersal can evolve to optimize trade-offs between finding habitats, avoiding kin competition, and colonizing new patches, as landscape patterns develop (Treep et al. 2020). Lifespan influences the ecological inheritance of niche construction (Odling-Smee et al. 2003). Ecological inheritance refers to the legacy of modified selection pressure by previous generations (Laland, Odling-Smee, and Feldman 1999). It posits that organisms not only transmit genes to subsequent generations, but also leave an environmental legacy that can affect selective pressures beyond their own lifetimes (Lehmann 2008). Such intertemporal phenotypic effects are also subject to natural selection (Lehmann 2008). However, constantly occurring landscape development will likely reduce the persistence of organismal environmental modifications and weaken the effect of ecological inheritance. For example, while plants can trap sediments to increase elevation, many physical processes modify the elevation in a complex way. When all the physical processes in the model are turned off, the evolution of niche construction is accelerated by >100% by forming large niche constructing spatial clusters in the phase of positive niche construction (Figs. 6 and S7).

Different mechanisms of niche construction, manifested over a broad range of environment-organism interactions, will vary in the likelihood that they will drive evolutionary responses (Matthews et al. 2014). Growth of sand dune grasses is a direct function of the amount of sand burial: plant growth is first enhanced by sand burial and then reduced by further burial in coastal sand dunes (Maun and Perumal 1999). This is the case of a direct relationship between environment-altering activities and modification of the selective environment. This applies to the model in this study. In other cases, the selective environment is not significantly modified by direct environment-altering activities, until specific geomorphic features are formed. For instance, weathering of limestone bedrock forms depressions, which retain more runoff and extend the inundation period for cypress trees growing within depressions (Dong, Murray, and Heffernan 2019). This benefits the survival of cypress trees, especially in water-limited systems. However, a homogeneous drop in elevation without forming an enclosed depression to hold runoff, would not significantly alter hydrological conditions—hence the selective environment— for cypress trees. While current niche construction theory suggests that across a broad range of conditions, niche construction traits can drive themselves to fixation (Silver and Di Paolo 2006) to generate unusual evolutionary dynamics (Laland, Odling-Smee, and Feldman 1999), results here imply that given the various sources of stochasticity, the likelihood of selection for niche construction traits might be limited, at least for plant niche construction in biogeomorphic landscapes.

### 4.3. Significance of coupling landscape development and biological evolution

Many plant niche construction traits, such as shoot density, root mass, and leaf shapes, have geomorphic consequences, but are not attributes specifically for niche construction (e.g., trapping sediments). They are instead multifunctional (Sack and Buckley 2020). For example, these traits could also facilitate photosynthesis, provide physical support, etc. The evolution of these traits would integrate tradeoffs among these functions (Sack and Buckley 2020). In this study, I assumed that traits exist exclusively for niche construction. In addition, I assumed that the species involved remains in the same system throughout landscape development. In reality, a given species often exists in a system during a certain stage of landscape development and then is replaced by one or more new species with better fit (i.e., they are fugitive species). This means that the species will experience intermittent positive niche construction phases, instead of the full phase of initial positive, later negative niche construction as depicted in this study (Fig. 3D). These properties provide a host of interesting research opportunities.

Many low gradient fluvial landscapes are biogeomorphic, as they have low stream power and vegetation can survive and flourish (Larsen 2019). As such, vegetation plays a much more profound role in these ecosystems than it does in others. Low gradient fluvial landscapes are globally significant in their fluxes and storage of water, sediment, and carbon, but are also vulnerable to climate change, e.g., loss of coastal marshes to sea level rise (Kirwan et al. 2016). Geomorphologists have increasingly realized complexities in the role of vegetation given the multiplicity of scales involved—from seconds to centuries to millennia, and from centimeters to 100s of kilometers (reviewed by (Larsen 2019))—many of which are relevant to evolution. Considering the simultaneous evolution of plants in these landscapes could have significant implications for climate change projections. For example, model results show that evolution of niche construction traits causes the landscape to stabilize at a higher elevation than it would otherwise (Fig. 2). This will influence the ability of coastal marshes to match accelerating sea level rise. While the implications of coupling evolution and landscape development are profound for low gradient fluvial landscapes, the conclusions in this paper are applicable to a wide range of systems where physical processes and vegetation interact. The term ‘evolution’ has been used commonly in geomorphology as in “landscape *evolution*.” This study highlights the need to distinguish biological evolution as a separate integral component necessary for understanding change in biogeomorphic landscapes. Likewise, evolutionary biologists need to recognize the significance of this landscape *evolution*, despite the fact that it does not involve changes in gene frequencies or natural selection.

