## Supplementary Materials for "Evolution of Plant Niche Construction Traits in Biogeomorphic Landscapes"

**Table S1.** Model parameters, their definitions, values used in the reference model.

| Parameter | Definition & dimension | Value in ref. model | Note |
|---|---|---|---|
| $Age_{max}$ | Maximum age of a plant [yr] | 30 | Three different levels were tested: 30, 400, 1500; 30 was used in the default model. |
| $Biom_{max}$ | Maximum biomass of a plant [-] | 1 | |
| $Biom_c$ | Threshold biomass above which the increase of seed production as a function of biomass slows down [-] | 0.5 | |
| $G_0$ | Parameter of the lognormal distribution for the pool of trait value of $G$ [-] | -7.8 | |
| $\sigma_G$ | Parameter of the lognormal distribution for the pool of trait value of $G$ [-] | 0.9 | |
| $R_0$ | Parameter of the lognormal distribution for the possible trait value of $R$ [m] | 1.5 | |
| $\sigma_R$ | Parameter of the lognormal distribution for the possible trait value of $R$ [m] | 0.18 | |
| $\beta_R$ | Constant to regulate the change in plant growth rate as a function of the difference between the actual elevation and the elevation preferred by plants [yr m$^{-2}$] | 10 | Three different levels were tested from low to high sensitivity: 0.4, 2, 10; 10 was used in the default model. |
| $\varepsilon$ | Shape parameter to describe the evolutionary cost of investing in 1 unit of niche construction trait $G$ [yr$^{-1}$] | 0.5 | |
| $\beta_m$ | Constant to regulate the change in plant mortality rate with the plant's age [yr$^{-1}$] | $\frac{0.01}{Age_{max}}$ | |
| $\alpha$ | Constant to regulate the effect of Euclidean distance on seed dispersal [m$^{-1}$] | 0.7 | Three different levels were tested from local to long-distanced dispersal: 0.7, 0.1, 0.01; 0.7 was used in the default model. |

| | | | |
|---|---|---|---|
| $p_s$ | Minimum probability for a plant older than age $A_s$ to produce seeds in a year [-] | 0.2 | |
| $A_s$ | The threshold age above which a plant can produce seeds [yr] | $0.1Age_{max}$ | |
| $\kappa$ | Shape parameter describing the rate of change in seed produced per unit of biomass [-] | 10 | |
| $b$ | Scale parameter to describe the effect of the mismatch between the actual and optimal elevation on successful seed germination [-] | 0.5 | |
| $\beta_G$ | Constant to describe the sediment trapping capacity of niche construction trait $G$ [m $yr^{-1}$] | 30 | Three different levels were tested from weak to strong niche construction capacity: 0.00001, 8, 30; 30 was used in the default model. |
| $d_1$ | Decay scaling factor of rate of elevational change [m] | 0.33 | |
| $d_2$ | Decay scaling factor of rate of elevational change [m] | 0.18 | |
| $k$ | Dimensionless coefficient ($k < 1$) that determines the magnitude of the rate when elevation is 0, relative to $(J + \varepsilon_G Biom_i G_i)$ [-] | 0.999 | |
| $D$ | Diffusion coefficient of sediments [m $yr^{-1}$] | 0.0005 | |
| $S_0$ | Negative effect received by a site by another site which is one meter higher and one-unit distance away [m $yr^{-1}$] | 0.0032 | |
| $\omega$ | Rate coefficient in the Gaussian function to describe the decaying of the negative effect as a function of distance [$m^{-2}$] | 0.25 | |
| $J$ | Baseline elevational change [m $yr^{-1}$] | 0.0008 | |

**Figure S1.** The effect of geo-evolutionary feedbacks, dispersal mode, and plant lifespan on correlation coefficient among the trait values of the geomorphic trait $G$ and the recipient trait $R$, and the elevation $h$. (A), (B), (C) show the effect of the strength of geo-evolutionary feedback ($L$-low: $\beta_R = 0.4, \beta_G = 0.00001$; $M$-medium: $\beta_R = 2, \beta_G = 8$; and $H$-high: $\beta_R = 10, \beta_G = 30$) on the steady-state, landscape-level mean correlation coefficient (the variation in the box plots captures the temporal variation over 100 kyrs) when dispersal is local and plant lifespan is short. (D), (E), and (F) show the effect of plant lifespan and dispersal mode on the landscape-level mean correlation coefficient over time under strong geo-evolutionary feedback. 95% confidence interval is created from 10 replicated simulations. Lifespan of 30 yrs was used for short-lived plants and 1,500 yrs was used for long-lived plants. Local dispersal, $\alpha = 0.7$ ($Eq.$ 4), and $\alpha = 0.01$ in long-distance dispersal.

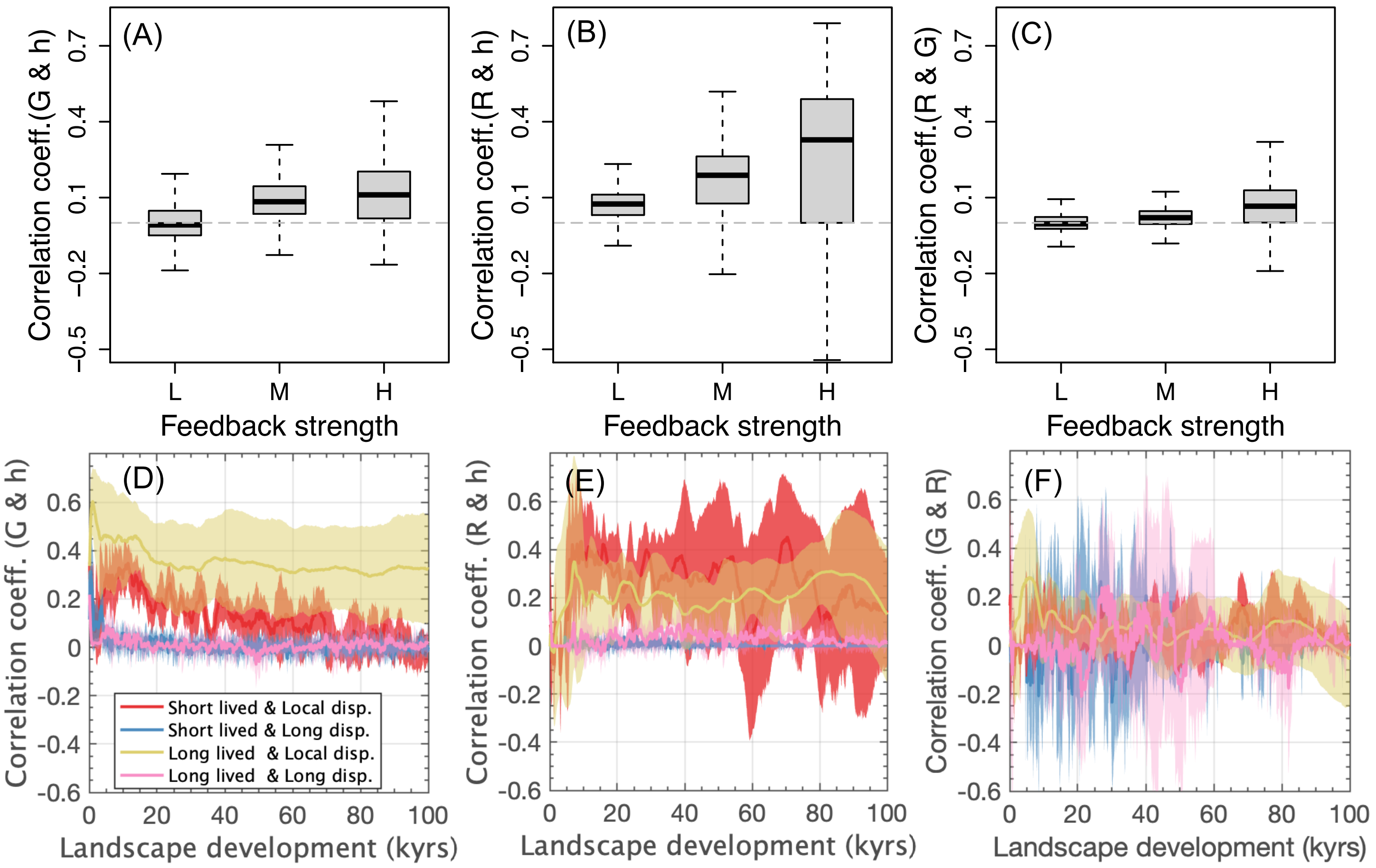

**Figure S2**. Correlation coefficient, diversity of *R* in the population, population mean value of the niche construction trait *G* in the model with no physical processes (A1-3), in the model with baseline elevational change only (B1-3), and in the reference model (C1-3). The time when the diversity of *R* peaks overlaps with the time of highest population mean *G* value and with the time of highest *G*-*R* correlation coefficient.

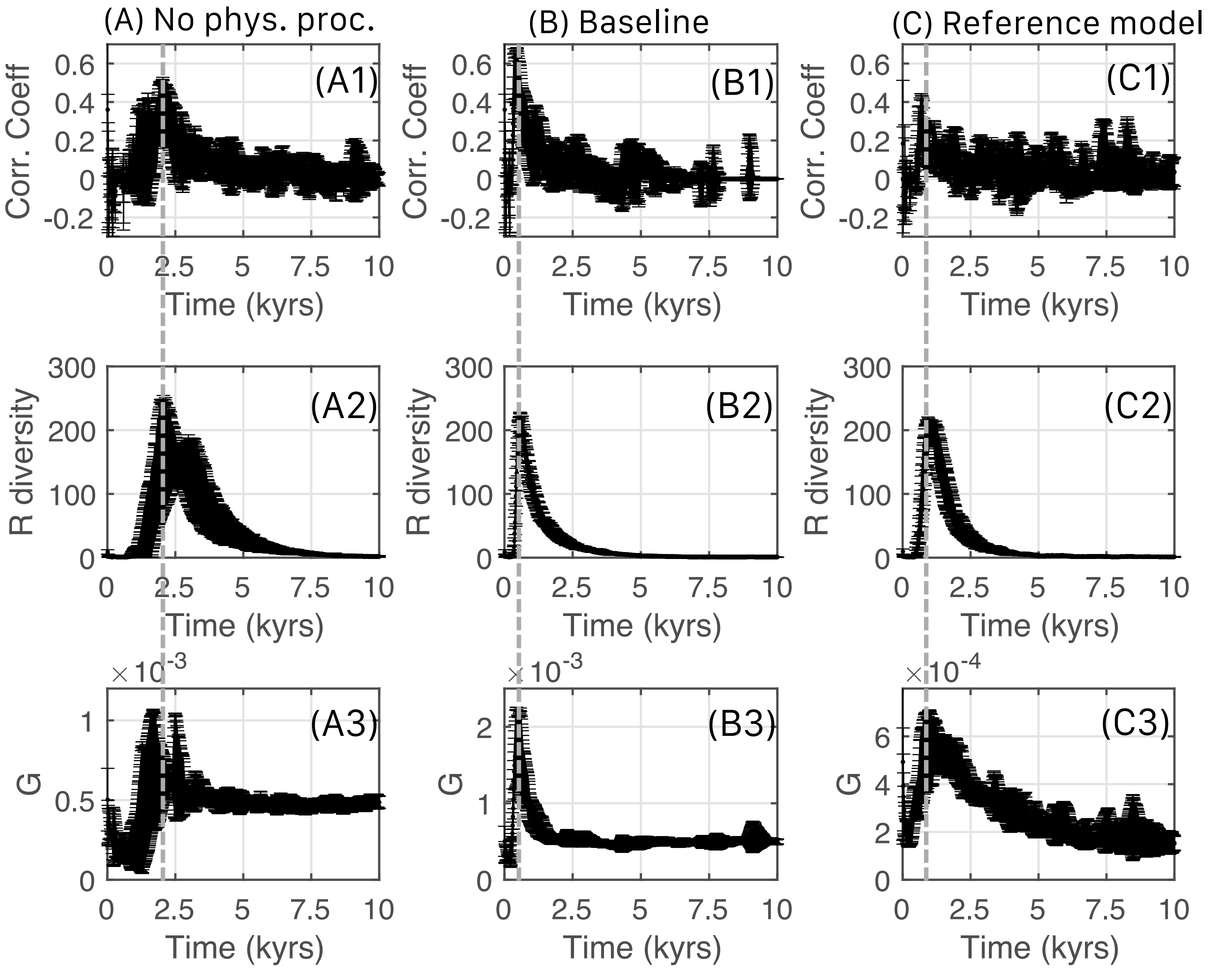

**Figure S3.** Landscape development, long-distance dispersal, and long lifespan increase stochasticity in the evolution of niche construction traits. "*SS*"–short-distance dispersal ($\alpha$ = 0.01) and short lifespan (30 yrs); "*MM*"–intermediate-distance dispersal ($\alpha$ = 0.1) and intermediate lifespan (400 yrs); "*LL*"–long-distance dispersal ($\alpha$ = 0.7) and long lifespan (1,500 yrs). Reference model has all the processes on; "*all off*" (B, E, H) has all the physical processes off, and the elevation in the model is only changed by plant niche construction; "*baseline*" (C, F, I) has all the physical processes, except baseline elevational change is off. Black open points connected by black lines are the population mean trait values averaged from 20 replicated simulations, with standard deviations (vertical red line for $R$ nd horizontal blue line for $G$).

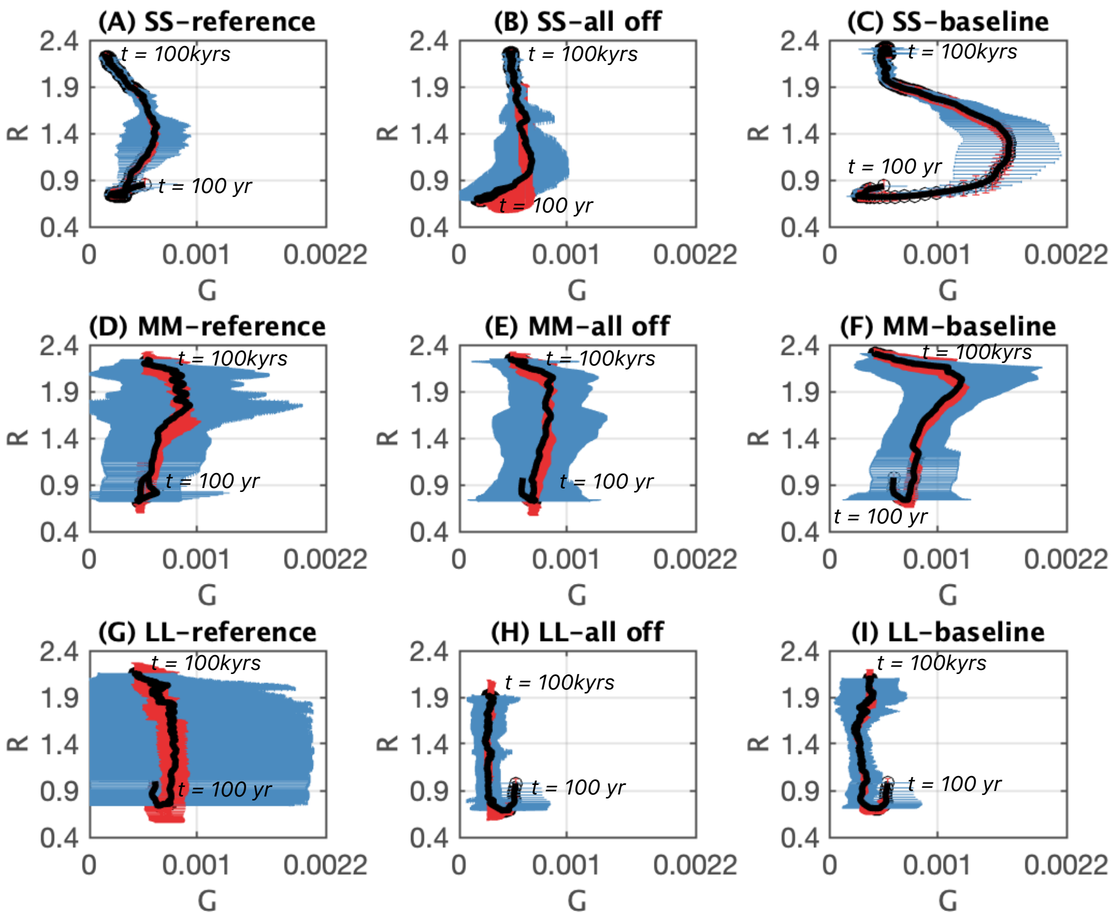

**Figure. S4**. Effect of turning on/off different physical processes of landscape development on the steady-state landscape pattern, with plant niche construction activities on. (A) All physical processes are turned off, and landscape develops by plant niche construction only; (B) Sediment diffusion is the only physical process turned on; (C) Sediment diffusion and long-distanced negative feedback are on; (D) Baseline elevational change is the only physical process on; (E) Sediment diffusion and baseline elevational change are turned on; and (F) all the three physical processes are turned on.

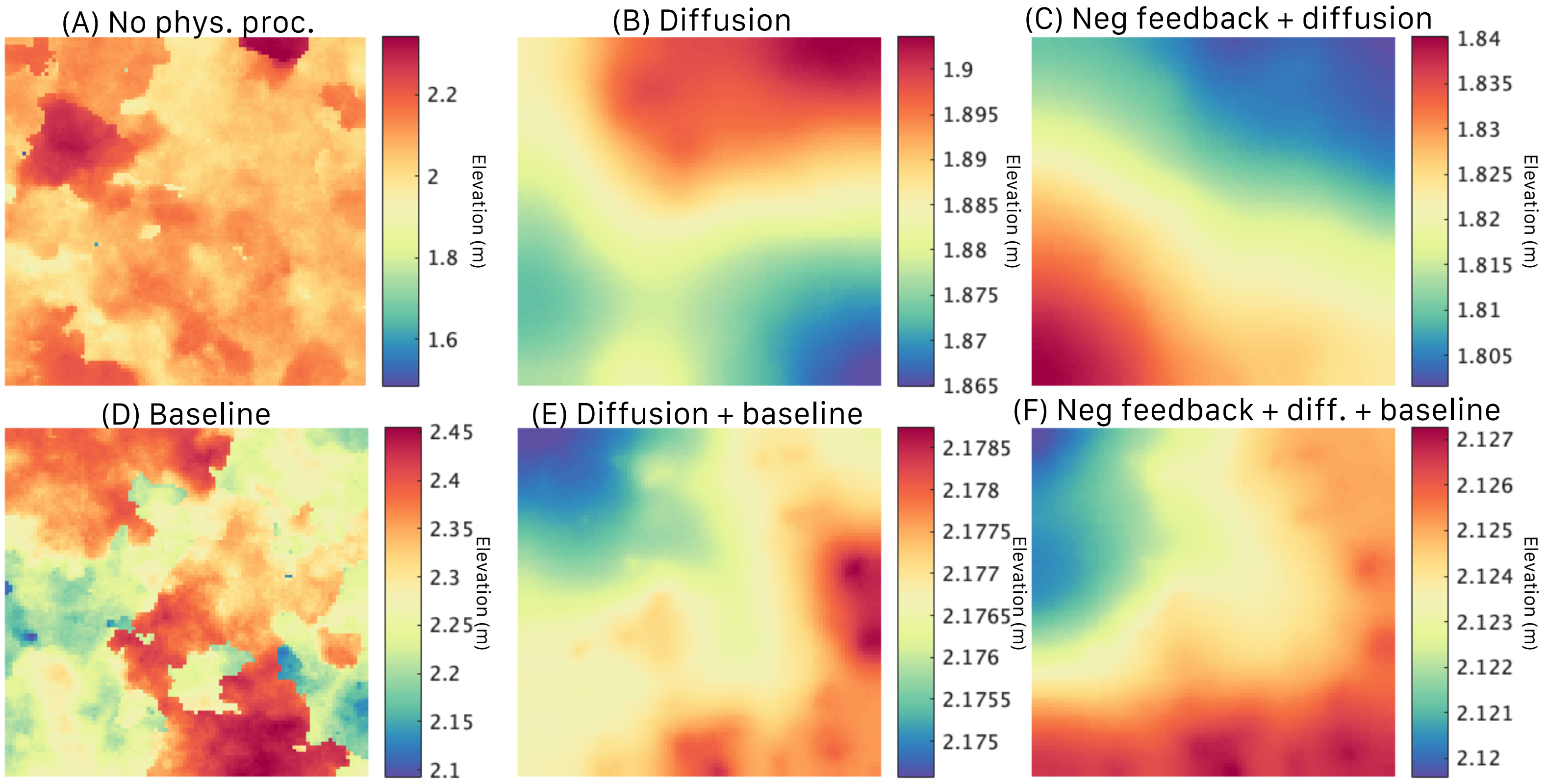

**Figure S5**. Dynamics of the recipient trait, *R*, and the geomorphic trait, *G***,** in the population and their expansion across the landscape (total size = 10,000 grids). Each hump-shaped curve represents the rise and fall of a mutation that lasted for at least 100 generations (~3,000 years). Each curve describes the time when the mutation rises, its abundance in the population over time, and the time when it disappears from the population.

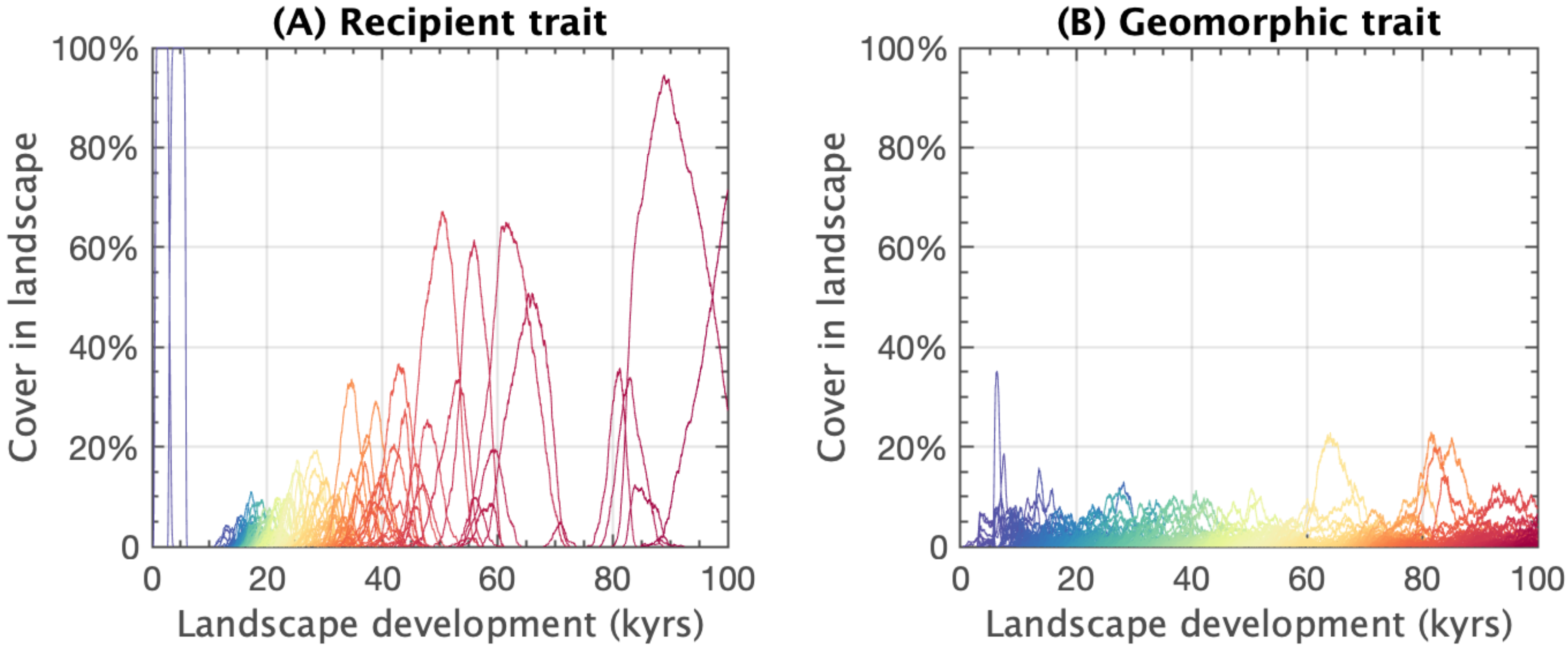

**Figure S6**. Spatial patterns of landscape elevation (A), biomass (B), trait value of recipient trait *R* (C), and trait value of the geomorphic trait *G* (D) in different stages of landscape development: 100 yrs, 10,000 yrs, 40,000 yrs, and 100,000 yrs. Each plot represents a model system with a domain of 100 x 100 grids.

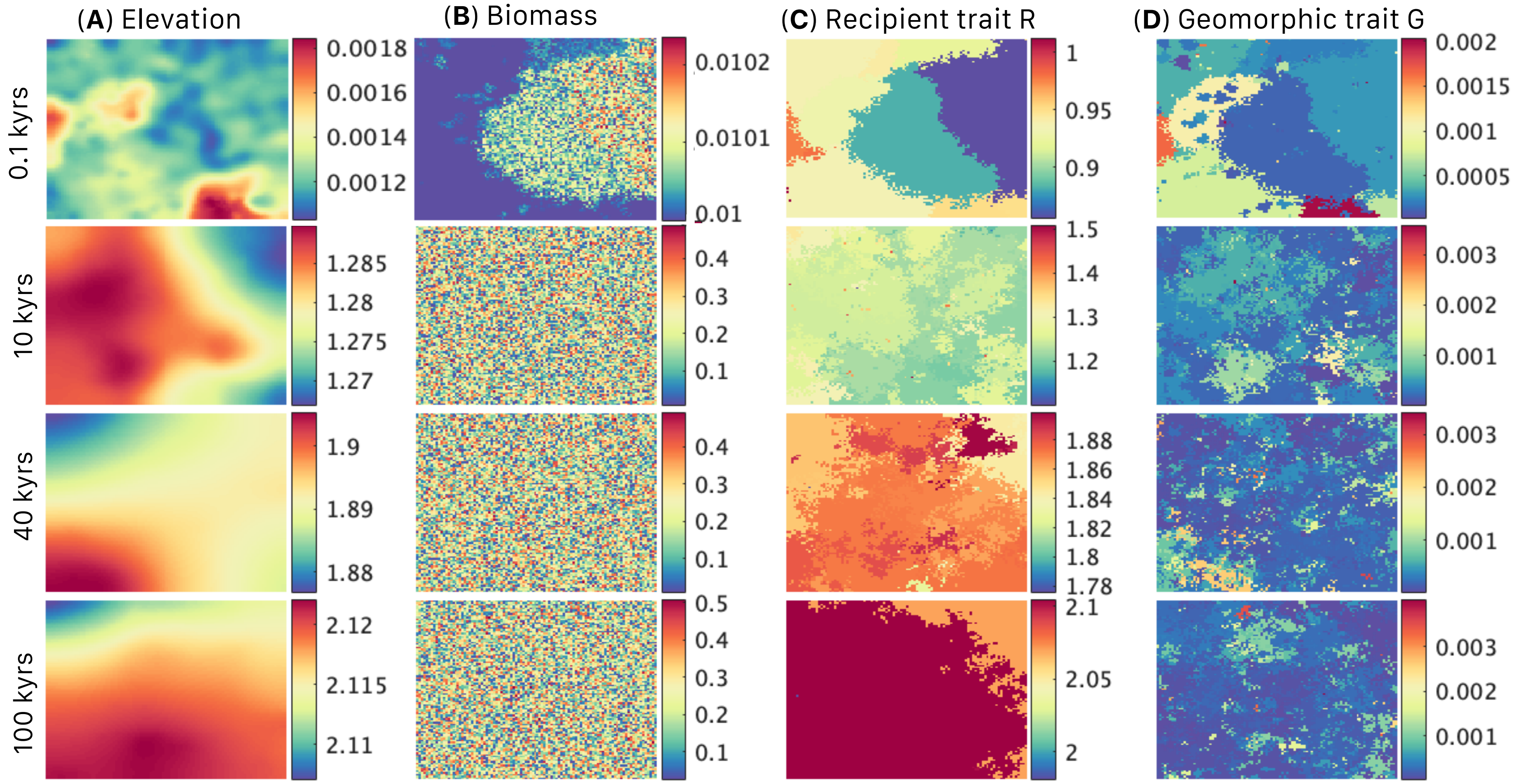

**Figure S7.** Differences in the evolutionary dynamics of the recipient trait $R$ and niche construction trait $G$, (A) when abiotic processes in landscape development are turned off (i.e., elevational change is solely caused by plant niche constructing activities), and (B) when landscape development occurs. The snapshots of elevation and trait patterns in (A) and (B) were taken from different periods in time—between 11–18 kyrs for the case when landscape development is turned off (A) and between 5–12 kyrs when landscape development is turned on, so that they have approximately similar elevations. In (A), the magenta-colored boxes highlight the development of a niche constructing spatial cluster: development of large-value $G$ (A17) increases the elevation in that location (A1), which makes large-value $R$ favored in the constructed environment (A9).

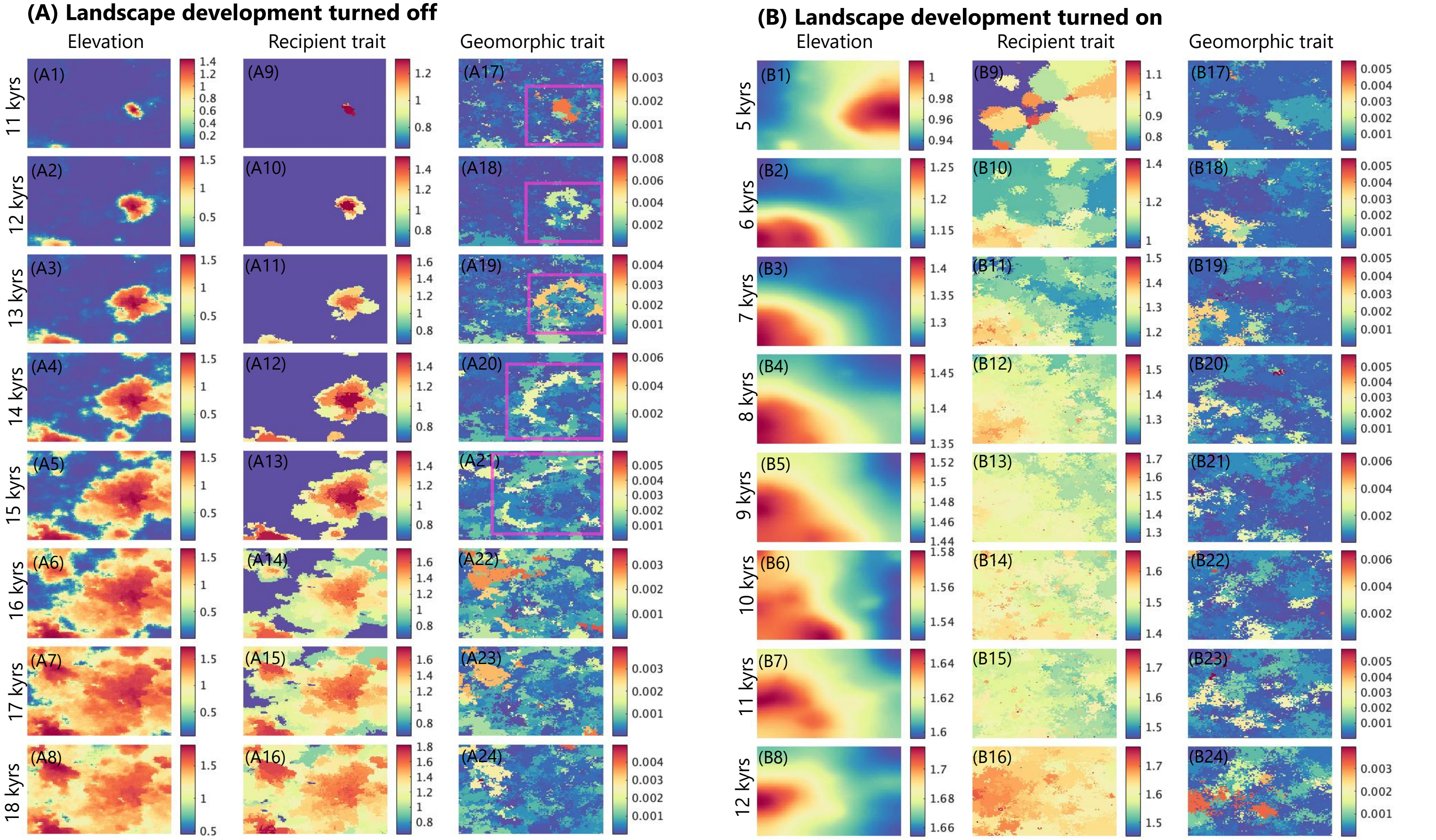

**Figure S8**. Increasing the evolutionary cost of niche construction trait *G* and increasing the rate of sediment diffusion make the evolution of *G* less likely to occur (greater cost of niche construction is represented by increasing $\varepsilon$ from 0.5 in the reference model to 50. Diffusion coefficient is increased from 0.0005 in the reference model to 0.003; the rest of the parameters are the same in the two models as given in Table S1)

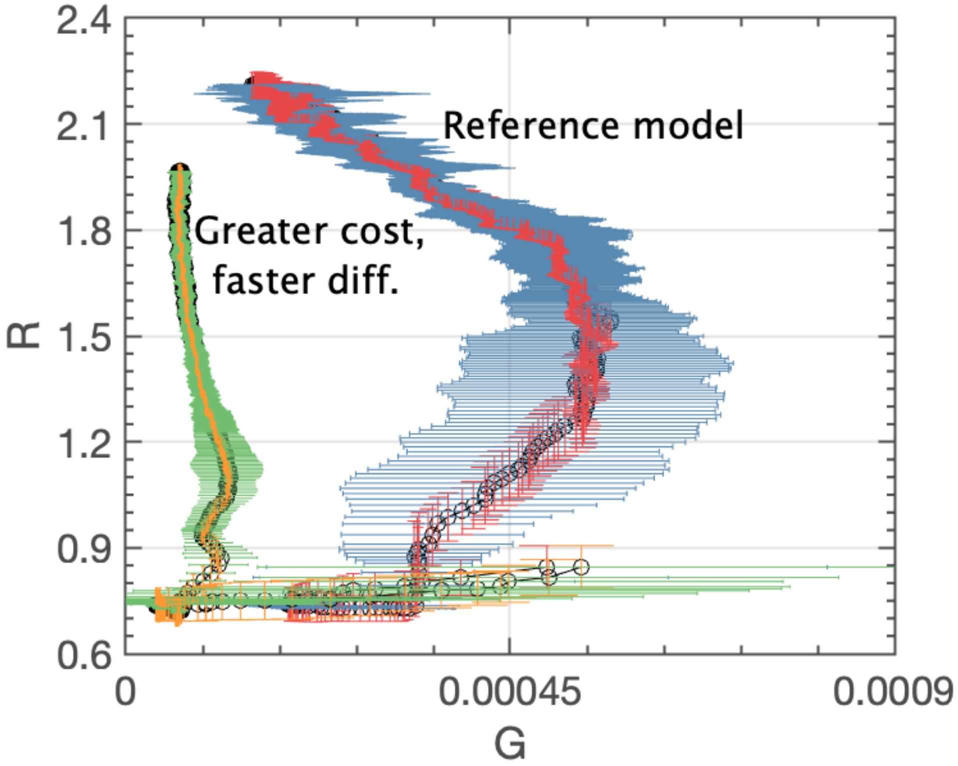
